# Periplasmic detoxification of urate hydroperoxide underpins *E. coli* survival in the inflamed gut

**DOI:** 10.64898/2026.06.01.729251

**Authors:** Maxence S. Vincent, Laurent Loiseau, Matthew Jones, Pierre Stocker, Khalid El Karkouri, Marion Chery, Divya Choudhary, Nicolas Vidal, Maria G. Winter, Natasha W. Tanner, Sylvia Pietri, Sebastian E. Winter, Benjamin Ezraty

## Abstract

Uric acid, the final product of purine metabolism in humans, accumulates in blood and tissues at relatively high concentrations^1^ as humans lack the enzyme uricase^2,3^. Under inflammatory conditions, uric acid can be oxidised to yield reactive intermediates^4^. In activated neutrophils, myeloperoxidase (MPO) catalyses the oxidation of uric acid by hydrogen peroxide, leading to the formation of urate hydroperoxide (UH)^5,6^. While recent studies have shown that UH is toxic to bacteria lacking peroxiredoxins^7^, its precise mechanism of toxicity and the existence of dedicated bacterial defence systems remain unknown. Here, we identify HiuH as a periplasmic enzyme, conserved across *E. coli* strains, that specifically degrades UH. Our findings reveal that UH selectively induces the expression of *hiuH* and that HiuH efficiently detoxifies UH both *in vitro* and in bacterial cells. HiuH cooperates with MsrP, a periplasmic methionine sulfoxide reductase that repairs UH-induced protein-bound methionine oxidation. This combined defence offering both direct detoxification and damage repair, is essential for bacterial survival under UH stress, and confers a competitive fitness advantage in a DSS-induced mouse model of colitis. Although UH is chemically transient, our work shows that it imposes durable biological consequences and a sufficient fitness cost in the in vivo niches occupied by *E. coli* to favour the evolution of a dedicated detoxification pathway beyond general oxidative-stress responses, defining a key adaptation to periods of gut inflammation.

## MAIN

Uric acid, which accumulates at unusually high concentrations in humans owing to the evolutionary loss of uricase, has long been considered a major circulating antioxidant. Increasing evidence, however, indicates that urate can also act as a pro-oxidant depending on the oxidative context^4^.

In particular, urate serves as a substrate for haem peroxidases such as MPO and lactoperoxidase that generate UH^6,8^, a recently identified reactive oxidant detected in activated human neutrophils^6^ and capable of modifying biomolecules^9–11^. During intestinal inflammation, neutrophil influx and peroxidase activity may favour urate oxidation within the inflamed gut environment, exposing luminal bacteria to UH. Yet how this short-lived oxidant affects bacterial physiology, colonisation and fitness remains largely unexplored.

In vitro, UH oxidises methionine residues^12^. In *E. coli*, reducing systems for oxidised methionine exist in both the cytoplasm and the periplasm^13^. The periplasmic system is encoded by the *msrPQ* genes, which form an operon with the upstream gene *hiuH*^14^ (**Fig.1a**). Expression of this operon is controlled by the HprSR two-component system upon oxidation of its periplasmic methionine^14^.

**Figure 1:**
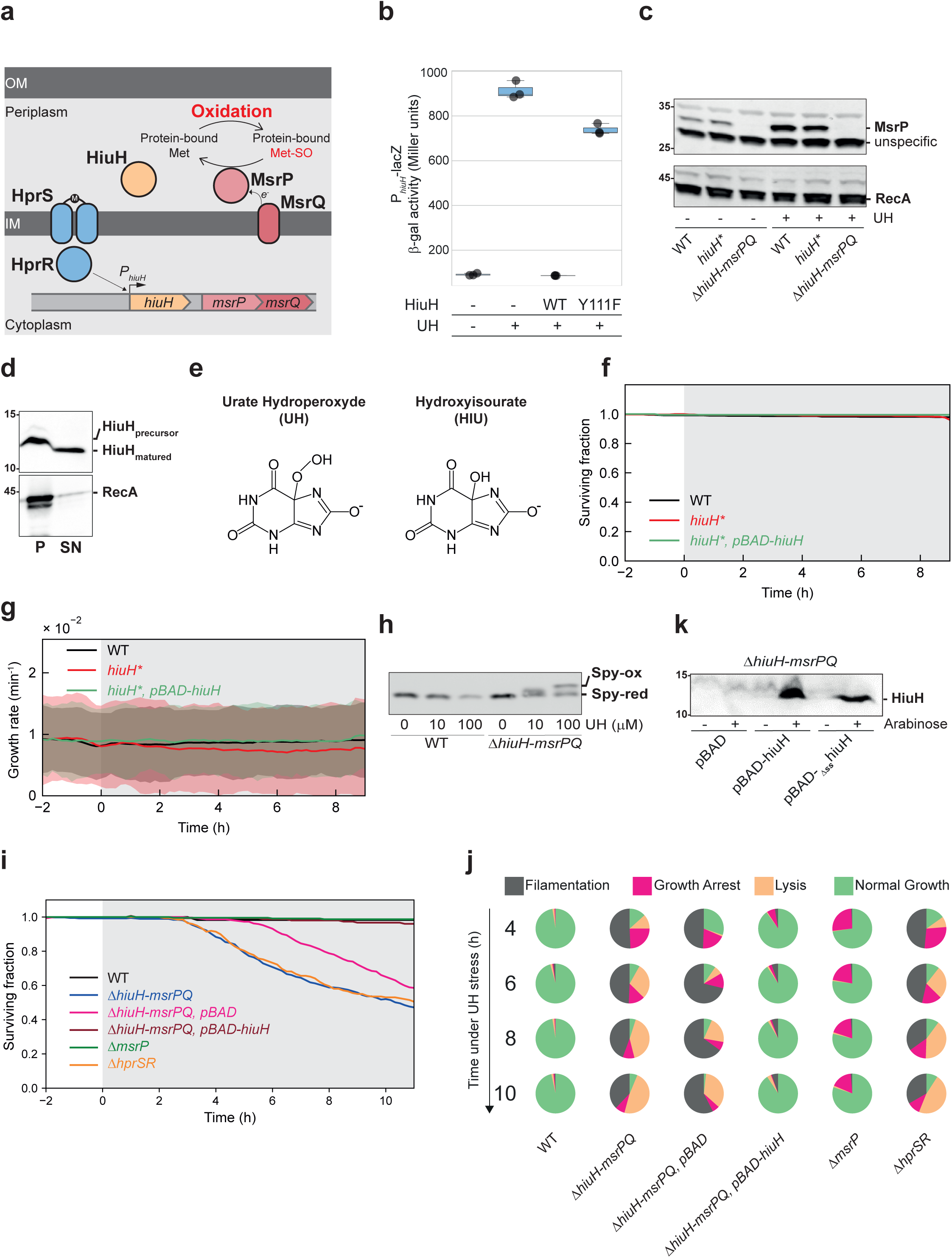
The *hiuH-msrPQ* operon is induced by UH and is essential for resistance to UH stress. **a.** Schematic of the *hiuH-msrPQ* operon and its regulation. *hiuH, msrP*, and *msrQ* are co-transcribed and regulated by the two-component system HprSR. OM, outer membrane; IM, inner membrane. b. UH induces expression of the *hiuH-msrPQ* operon. β-galactosidase activity from a chromosomally integrated P*_hiuH_*-*lacZ* transcriptional fusion (ectopic locus) was measured in cells exposed to: (i) no UH, (ii) UH alone, (iii) UH pre-incubated with purified HiuH, or (iv) UH pre-incubated with the inactive HiuH^Y111F^ variant. Box plots show data from three independent biological replicates; individual values are overlaid. c. UH exposure increases MsrP protein levels. Total cell extracts from WT, *hiuH*,* and *ΔhiuH-msrPQ* strains were analysed by immunoblotting using anti-MsrP antibodies, with or without UH treatment. Anti-RecA was used as a loading control. Molecular weight markers (kDa) are indicated. d. HiuH is periplasmic protein. Western blot analysis with anti-His antibodies was performed on fractionated extracts from a strain expressing HiuH^6His^. Anti-RecA was used to control for cytoplasmic contamination. e. Chemical structures of urate hydroperoxide (UH) and hydroxyisourate (HIU). f. Survival ratio during constant UH exposure (shaded area) for WT (n=585), *hiuH\** (n=388), and *hiuH\** carrying pBAD-*hiuH^6His^* (n=208). g. Population-averaged growth rates under UH exposure. Bold lines represent the mean, and shaded areas indicate the standard deviation across single-cells. h. HiuH-MsrPQ prevent UH-oxidation of the periplasmic chaperone Spy. WT and *ΔhiuH-msrPQ* strains were exposed to increasing concentrations of UH, and total cell extracts were analysed by immunoblotting using anti-Spy antibodies. The oxidation state of Spy, reflected by its electrophoretic mobility shift, is indicated. i. Survival ratio during constant UH exposure (shaded area) for WT (n=116), *ΔhiuH-msrPQ* (n=369), *ΔhiuH-msrPQ* carrying an empty pBAD (n=151), *ΔhiuH-msrPQ* carrying pBAD-*hiuH^6his^* (n=292), *ΔmsrP* (n=226) and *ΔhprSR* (n=198). j. Quantification of physiological state fractions over time for the same strains and conditions as in panel i. k. Western blot analysis with anti-His antibodies on total extracts from the *ΔhiuH-msrPQ* strain carrying an empty pBAD, *pBAD-hiuH^6His^*, or *pBAD-_Δss_hiuH^6His^* (signal sequence deleted), with or without arabinose induction. Molecular weight markers (kDa) are indicated.

### The *hiuH-msrPQ* operon is essential during UH exposure

UH can be generated through the photoactivation of riboflavin, a reaction that produces superoxide anions which subsequently react with urate radicals to yield UH^15,16^. Building on established photooxidative protocols^15,17^, we adapted this system to reproducibly generate UH under controlled conditions (**Extended Data Fig.1a,b,c,d**), and found that UH exposure induces expression of the *hiuH-msrPQ* operon (**Fig.1b,c and Extended Data Fig.1e,f,g**).

While the role of MsrPQ in repairing oxidised methionine is well established^18^, the function of HiuH is less understood. HiuH is an orthologue of PucM, a hydroxyisourate (HIU) hydrolase from *Bacillus subtilis*^19^, whose physiological significance in *E. coli* remains unclear because (i) HIU is produced in the cytoplasm, whereas HiuH localises in the periplasm (**Fig.1d**), (ii) *E. coli* lacks uricase, the enzyme that produces HIU during purine catabolism. However, UH and HIU exhibit very similar chemical structures (**Fig.1e**), which led us to hypothesise that HiuH could work on UH instead of HIU.

To specifically impair HiuH function without disrupting the downstream operon, we introduced a premature stop codon into *hiuH*, generating the *hiuH\** strain. MsrP was similarly upregulated upon UH exposure in WT and *hiuH\** cells, indicating that operon induction was preserved (**Fig.1c**).

We assessed the contribution of HiuH to UH tolerance at the single-cell level using a light-inducible microfluidic system in which UH is generated in situ from riboflavin and uric acid in a defined M9-based growth medium, while bacteria are monitored over time within a mother-machine device (**Extended Data Fig.2a,b,c**). Under these conditions, *hiuH\** cells showed no detectable survival defect compared with WT cells (**Fig.1f**), but displayed a reduced growth rate (**Fig.1g**), suggesting that HiuH contributes to maintaining cellular fitness during UH exposure.

We thus wondered whether the *hiuH-msrPQ* operon could act as a dual defence mechanism in which HiuH would degrade UH directly, while MsrPQ would repair UH-induced damage. Consistent with this hypothesis, we found that UH oxidises the periplasmic chaperone Spy, a known MsrPQ client^20^, and that deletion of the operon abolished repair of this damage (**Fig.1h**). More importantly, the loss of the operon severely compromised survival under UH stress, but the complementation of the mutant with *hiuH* alone was sufficient to restore significant viability (**Fig.1i,j,k and Supplementary Video 1**), indicating that overproduction of HiuH can compensate the loss of MsrPQ.

Of note, survival of Δ*hiuH-msrPQ* cells was impaired only when both riboflavin and uric acid were present, indicating that toxicity was UH-dependent and not caused by reactive species generated from either excited compound alone (**Extended Data Fig.3**). We further confirmed that direct detoxification by HiuH is sufficient to prevent lethal damage by measuring survival levels that were unaffected by UH stress in the Δ*msrP* strain (**Fig.1i,j**). By contrast, the Δ*hprRS* mutant, which is unable to induce the *hiuH-msrPQ* operon, exhibited a complete loss of viability (**Fig.1i,j**).

### UH causes a periplasmic stress

When exposed to UH stress, most Δ*hiuH-msrPQ* cells ceased growth and eventually lysed, as evidenced by a sudden loss of cytoplasmic content observed during live-cell imaging (**Fig.2a,b and Supplementary Video 1**). Given that UH is a charged molecule, we suspected that cellular damage might initially occur in the periplasm, as UH would likely not passively diffuse across the inner membrane. Relying on multiple fluorescent compartment markers, we visualised the subcellular progression of UH-induced damage in real-time (**Supplementary Video 2**). In the *ΔhiuH-msrPQ* mutant, we observed a rapid release of the periplasmic content followed by the release of the cytoplasmic material (**Fig.2c,d,e**). This suggests that UH-induced damage initiates in the periplasm and subsequently affects the cytoplasm. Consistently, membrane damage was more pronounced in the *ΔhiuH-msrPQ* mutant than in the WT (**Fig.2d and Supplementary Video 3**), indicating that membrane integrity is compromised without the HiuH-MsrPQ periplasmic system.

**Figure 2:**
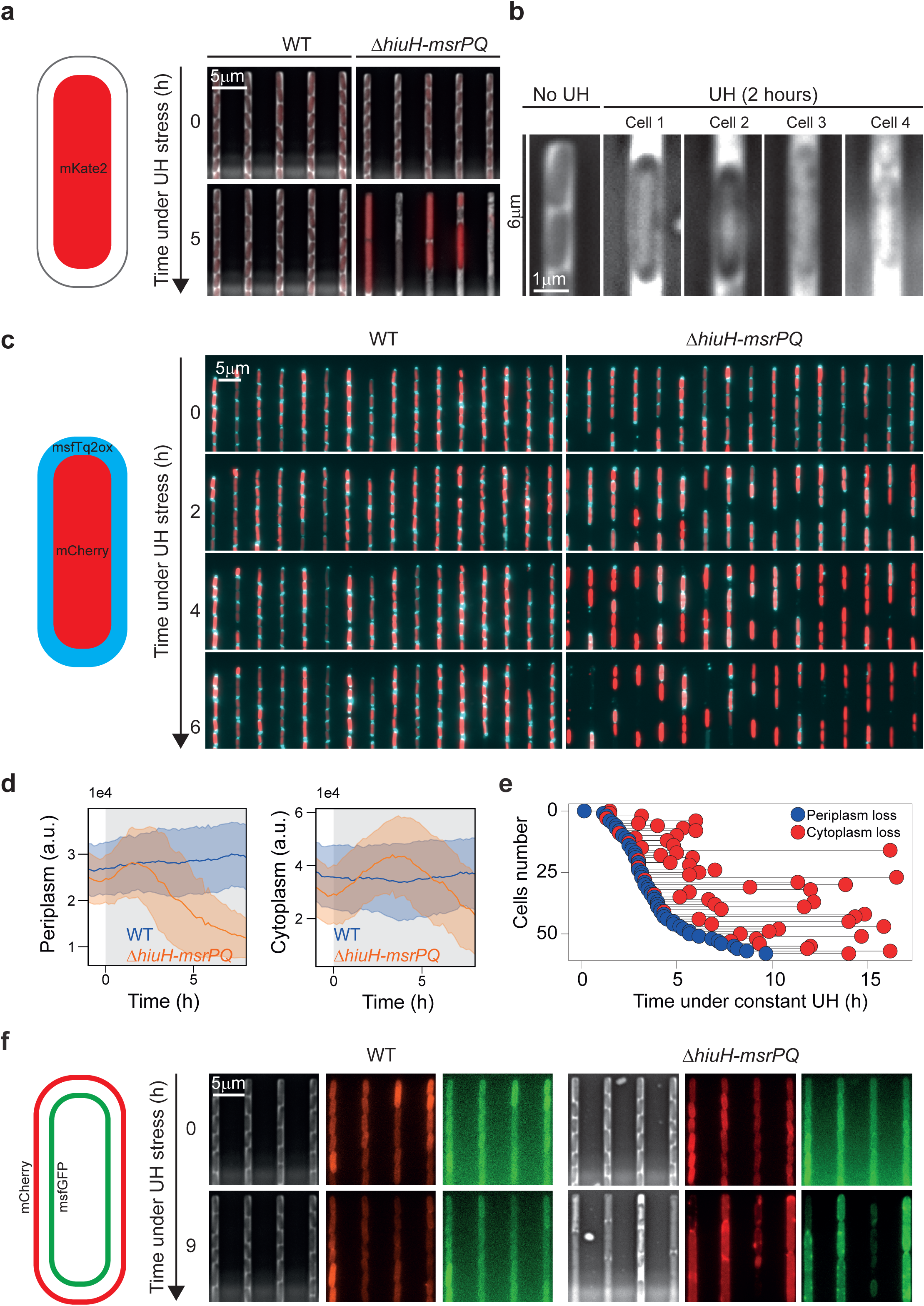
UH stress causes periplasmic damage and disrupts envelope integrity. **a.** Snapshots of WT (left) and *ΔhiuH-msrP* (right) strains expressing a chromosomally encoded constitutive cytoplasmic red fluorescent marker (mKate2), imaged in a single-cell microfluidic device before (0 h) and after 5 h of constant UH treatment. Phase contrast and red fluorescence are merged and normalised across all displayed images. Scale bar, 5 µm. b. Example phase-contrast images of *ΔhiuH-msrP* cells before UH treatment (left) and four representative *ΔhiuH-msrP* cells after ≥2 h of UH exposure (right). Cells are shown in phase contrast only. c. Wt (left) and *ΔhiuH-msrP* (right) strains expressing a plasmid-borne cytoplasmic red marker (mCherry) and a periplasmic blue marker (msfTq2ox), taken at 0, 2, 4, and 6 h of constant UH exposure. Scale bar, 5 µm. d. Time-resolved quantification of the population-averaged periplasmic (left) and cytoplasmic (right) fluorescence intensities in WT and *ΔhiuH-ΔmsrP* strains during UH treatment (shaded area). e. Distribution of cytoplasmic and periplasm release time for 63 *ΔhiuH-msrP* single cells. The boxplot in the inlet shows the delay between periplasm and cytoplasm loss for each single-cell. f. Snapshots of WT (left) and *ΔhiuH-msrP* (right) strains expressing an outer membrane marker (LpoB-mCherry) and an inner membrane marker (msfGFP-GlpT), imaged before and after 10 h of constant UH treatment. Scale bar, 5 µm.

To assess the importance of HiuH localisation, we expressed a cytoplasmic variant of the protein (_Δss_HiuH) in the *ΔhiuH-msrPQ* strain (**Fig1.k and Extended Data Fig.4a**) and found that it failed to restore growth under UH exposure (**Supplementary Video 4**). However, in contrast to the *ΔhiuH-msrPQ* mutant, which undergoes extensive filamentation prior to lysis, cells expressing cytoplasmic HiuH essentially exhibited growth arrest, with minimal filamentation before death (**Extended Data Fig.4a,c**). While this observation could suggest a protective effect of HiuH in the cytoplasm, it underscores the critical requirement of its periplasmic localisation to ensure normal growth during UH stress. In this regard, when the complete cytoplasmic methionine sulfoxide reduction system (MsrA, MsrB, MsrC, BisC) was disrupted, cell survival remained unaffected (**Extended Data Fig.4b,c**). This result reinforces that the periplasmic HiuH-MsrPQ system alone is sufficient to prevent UH from reaching the cytoplasm.

### Detoxification of UH depends on HiuH

Our microfluidic system enables real-time visualisation of the *hiuH-msrPQ* operon dynamics at single-cell resolution. Using a fluorescent reporter for *hiuH* promoter activity (*P_hiuH_-mYPet*), we measured that the operon is induced rapidly, within 10 minutes of UH exposure (**Fig.3a**). Interestingly, the response amplitude varied markedly among isogenic cells (**Fig.3b,c**). Previous studies have shown that clonal *E. coli* populations exhibit heterogeneous responses to H_2_O_2_, partly due to cell–cell protective effects^21^. Peripheral “barrier” cells scavenge H_2_O_2_, generating spatial gradients that reduce stress for adjacent neighbours^21^. We found that a similar phenomenon occurs under UH stress. Indeed, reporter expression revealed a gradient of response magnitude along the trenches of the microfluidic chip (**Fig.3d and Supplementary Video 5**). Upon UH exposure, cells at the open end of the trenches displayed stronger reporter activation, with signal intensity progressively decreasing towards the closed end (**Fig.3e**). Importantly, this gradient was UH-specific, as the *P_hiuH_-mYPet* reporter was selectively activated by UH but not by H_2_O_2_ (**Fig.3f**). By training a machine learning model on our dataset, we confirmed that collective cell-cell protection drives the observed gradient effect, with barrier cells as key predictors of the UH stress response heterogeneity (**Extended Data Fig.5**).

**Figure 3:**
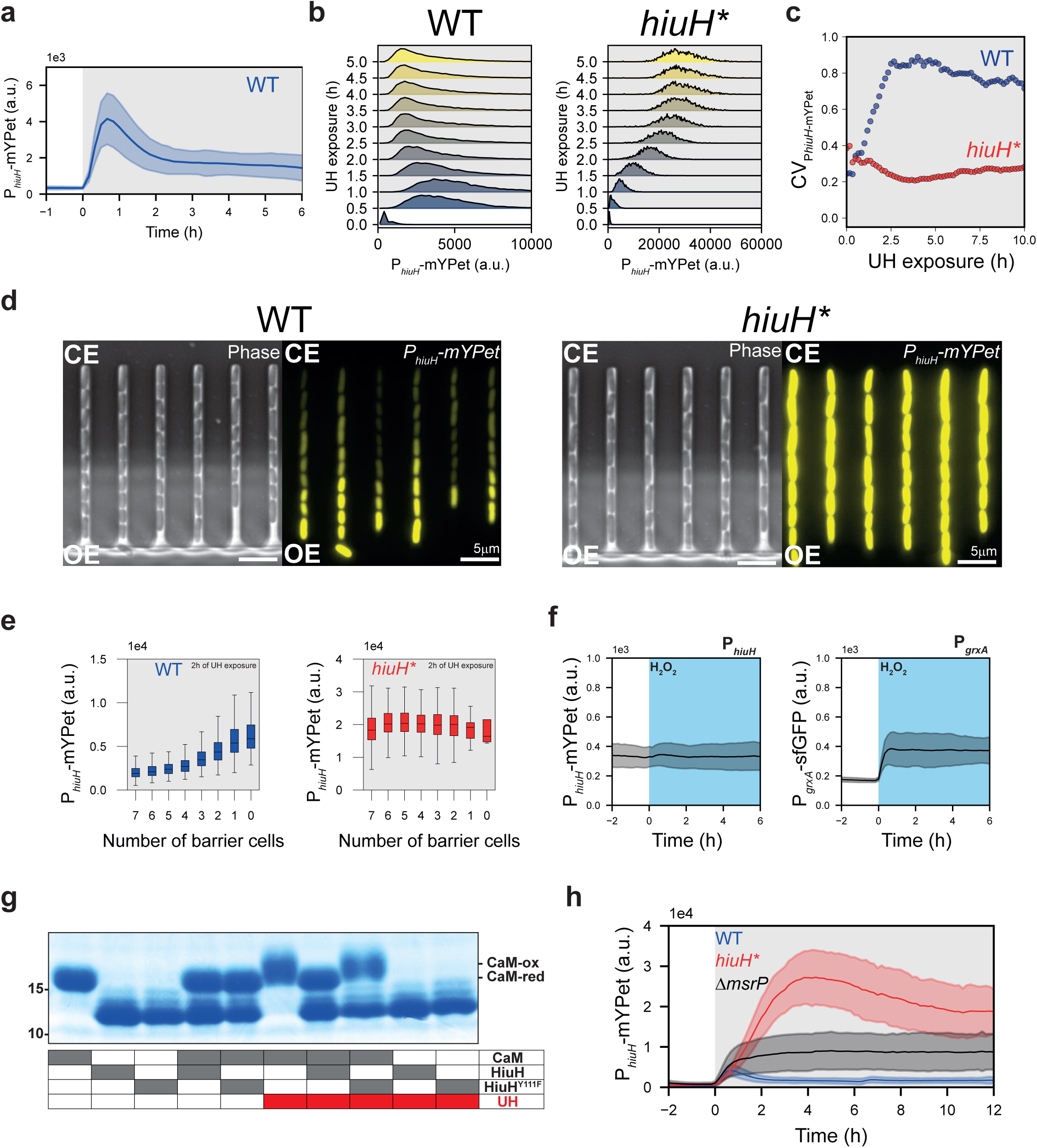
HiuH detoxifies UH. **a.** Time-resolved population-averaged expression of P*_hiuH_*-mYPet under constant UH treatment (shaded area). The blue bold line represents the mean; the blue shaded area indicates the standard deviation. B. Single-cell distributions of P*_hiuH_*-mYPet fluorescence levels in WT (left) and *hiuH* (*right) strains at 0, 0.5, 1, 1.5, 2, and 2.5 h after constant UH exposure. c. Coefficient of variation of P*_hiuH_*-mYPet fluorescence across single cells during UH treatment in WT (blue) and *hiuH*,* (orange) strains. d. Snapshots showing P*_hiuH_*-mYPet expression after 2 h of constant UH exposure in WT (left) and *hiuH\** (right) strains. Phase contrast and mYPet fluorescence channels are shown. CE, closed end; OE, open end of the microfluidic channel. Scale bar, 5 µm. e. Box plots showing P*_hiuH_*-mYPet fluorescence levels after 2 h of UH treatment in WT and *hiuH* st*rains, plotted as a function of the number of barrier cells. f. Time-resolved population-averaged fluorescence levels of P*_hiuH_*-mYPet (left) and P*_grxA_*-sfGFP (right) under constant H_2_O_2_ treatment (shaded area). g. Coomassie blue-stained SDS-PAGE of purified calmodulin (CaM) incubated with or without UH, in the presence or absence of purified HiuH or the catalytically inactive HiuH^Y111F^ variant. The redox states of CaM are indicated. Molecular weight markers (kDa) are shown. h. Time-resolved population-averaged P*_hiuH_*-mYPet expression under constant UH treatment in WT (blue), *hiuH\** (red), and *ΔmsrP* (black) strains. The bold lines represent the mean; the shaded areas indicate the standard deviation.

We reasoned that if neighbouring cells protect those behind them by activating a putative UH detoxification system, disrupting this system should abolish the gradient. We thus monitored gradient formation in the *hiuH\** mutant strain and found that the gradient was indeed completely abolished (**Fig.3b,c,d,e and Supplementary Video 5**). This result indicates that HiuH actively reduces UH availability in live bacterial populations. In agreement with this scenario, incubating HiuH with UH was sufficient to inhibit UH-induced activation of the *hiuH-msrPQ* operon (**Fig.1b**) and effectively impaired protein-bound methionine oxidation (**Fig.3g**).

Furthermore, the dynamics of P*_hiuH_* expression in the *hiuH*\* mutant (**Fig.3h**) supports a negative feedback loop regulating the *hiuH-msrPQ* operon, as previously proposed¹⁰. In the WT strain, P*_hiuH_* reporter activity decreases rapidly following UH exposure, consistent with HiuH-mediated UH degradation and consequent inactivation of the operon (**Fig. 3a**). By contrast, the *hiuH*\* mutant not only exhibits a much slower decline but also maintains a P*_hiuH_* signal nearly tenfold higher than WT, reflecting the inability to degrade UH (**Fig.3h**). The residual decrease in the *hiuH*\* mutant likely stems from MsrP activity, which reduces HprS activation by repairing its UH-oxidised methionine residues. Supporting this interpretation, a *ΔmsrP* strain shows a reduced amplitude of P*_hiuH_* induction compared to *hiuH*\*, but maintains reporter expression at a steady-state level, suggesting that in the absence of MsrP, the response cannot be efficiently switched off (**Fig.3h**).

### HiuH specifically degrades UH

To characterise the enzymatic activity of HiuH, we assessed its ability to degrade UH in vitro. Incubation of UH with purified HiuH led to a significant increase in degradation rate compared to spontaneous decay (**Fig.4a,b**), indicating direct catalytic activity. We next compared HiuH to its *Bacillus subtilis* orthologue PucM. Although both enzymes degraded HIU with comparable efficiency (**Fig. 4c,d**), HiuH was more effective than PucM at degrading UH, particularly at equivalent enzyme concentration (**Fig. 4a,b**). In line with this difference in activity, in vivo, periplasmic expression of PucM failed to restore UH resistance in the *ΔhiuH-msrPQ* mutant (**Fig. 4e and Supplementary Video 6**). These results reveal a functional divergence between HiuH and PucM, suggesting that HiuH has evolved enhanced specificity for UH despite its high structural similarity to PucM (**Fig.4f**).

**Figure 4:**
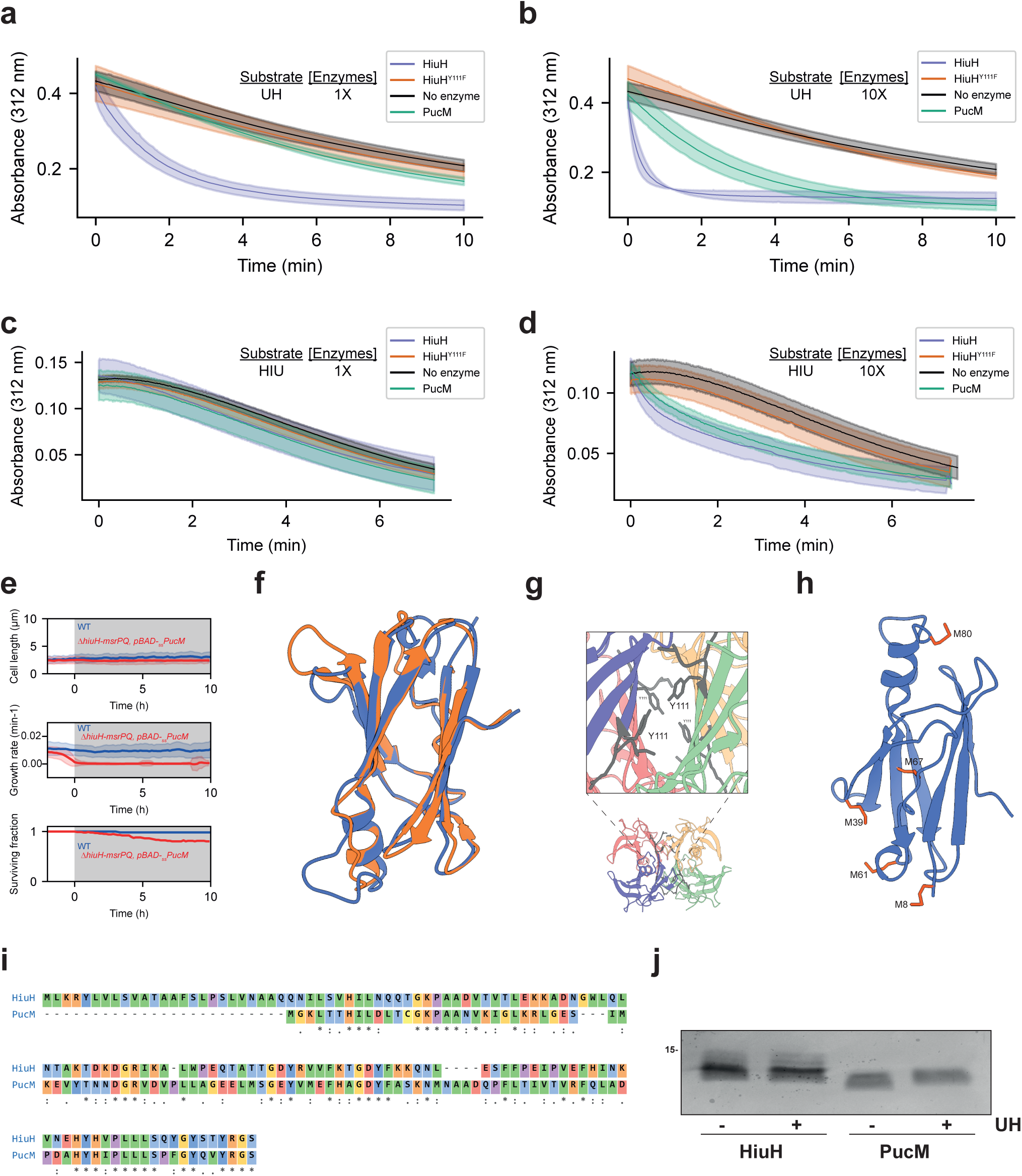
Functional divergence of HiuH and PucM underlies adaptation to UH detoxification. **a-d.** Degradation of UH (a, b) or HIU (c, d) over time in the absence or presence of HiuH, HiuH^Y111F^, or PucM. Enzymes were used at 0.06 μM (a, c) or 0.6 μM (b, d) (1X and 10X, respectively). Data represent the mean ± s.d. of at least three independent replicates. e. Population-averaged time-traces of cell length (top), growth rate (middle), and survival (bottom) for WT and *qhiuH-msrPQ* complemented with a pBAD plasmid expressing *pucM*, under continuous exposure to UH (grey shaded area). f. Superposition of HiuH (PDB: 2G2N) and PucM (PDB: 2H0E) crystal structures. g. Structure of the HiuH tetramer with a close-up of the catalytic site; Y111 residues from each monomer are indicated. h. Structure of PucM with methionine residues highlighted in orange. i. CLUSTALW-generated sequence alignment between HiuH (sp|P76341|HIUH_ECOLI) and PucM (sp|O32142|HIUH_BACSU). j Anti-His western blot showing the migration of purified HiuH^6His^ and PucM^6His^ with or without incubation with UH. Molecular weight is indicated.

The catalytic site of PucM is well characterised and fully conserved in HiuH (**Fig.4g**). Mutation of a key active-site residue (Y111F) in HiuH abolished UH-degrading activity in vitro (**Fig.4a**). In line with this, pre-incubation of UH with the purified HiuH^Y111F^ variant failed to inhibit UH-induced activation of the *hiuH-msrPQ* operon (**Fig.1b**) and did not protect against protein-bound methionine oxidation (**Fig.3g**), unlike WT HiuH. Because the catalytic residue required for HIU degradation by PucM is also essential for UH degradation by HiuH, and given that both enzymes retain activity toward HIU, we reasoned that the difference in UH degradation likely does not stem from divergent catalytic mechanisms.

Instead, we found that HiuH is completely devoid of methionine residues, whereas PucM contains five (**Fig. 4h,i**). Upon UH exposure, PucM underwent oxidation, while HiuH remained unaffected (**Fig. 4j**), suggesting that methionine oxidation may underlie the loss of PucM activity. Together, these findings indicate that HiuH is both catalytically and structurally adapted to degrade UH under oxidative conditions.

### The HiuH-MsrPQ module is conserved across *E. coli* strains

To place HiuH in an evolutionary context and distinguish it from canonical PucM-like HIU hydrolases, we surveyed bacterial proteins containing a hydroxyisourate hydrolase domain using sequence similarity, predicted signal peptide, subcellular localisation, methionine content and genomic context (**Extended Data Fig.6 and Table S1**). This analysis identified a restricted group of HiuH-like proteins predicted to be exported to the periplasm and to contain few or no methionine residues in the mature protein.

In *E. coli*, HiuH homologues were highly conserved and embedded in the same *hprS-hprR-hiuH-msrP-msrQ* genomic architecture, indicating that regulatory sensing, UH detoxification and methionine-sulfoxide repair are genetically linked in this species (**Fig.5a**). By contrast, more distant HiuH/PucM-like homologues were frequently found in variable genomic contexts and were not consistently associated with msrPQ (**Extended Data Fig.6 and Table S2**).

**Figure 5:**
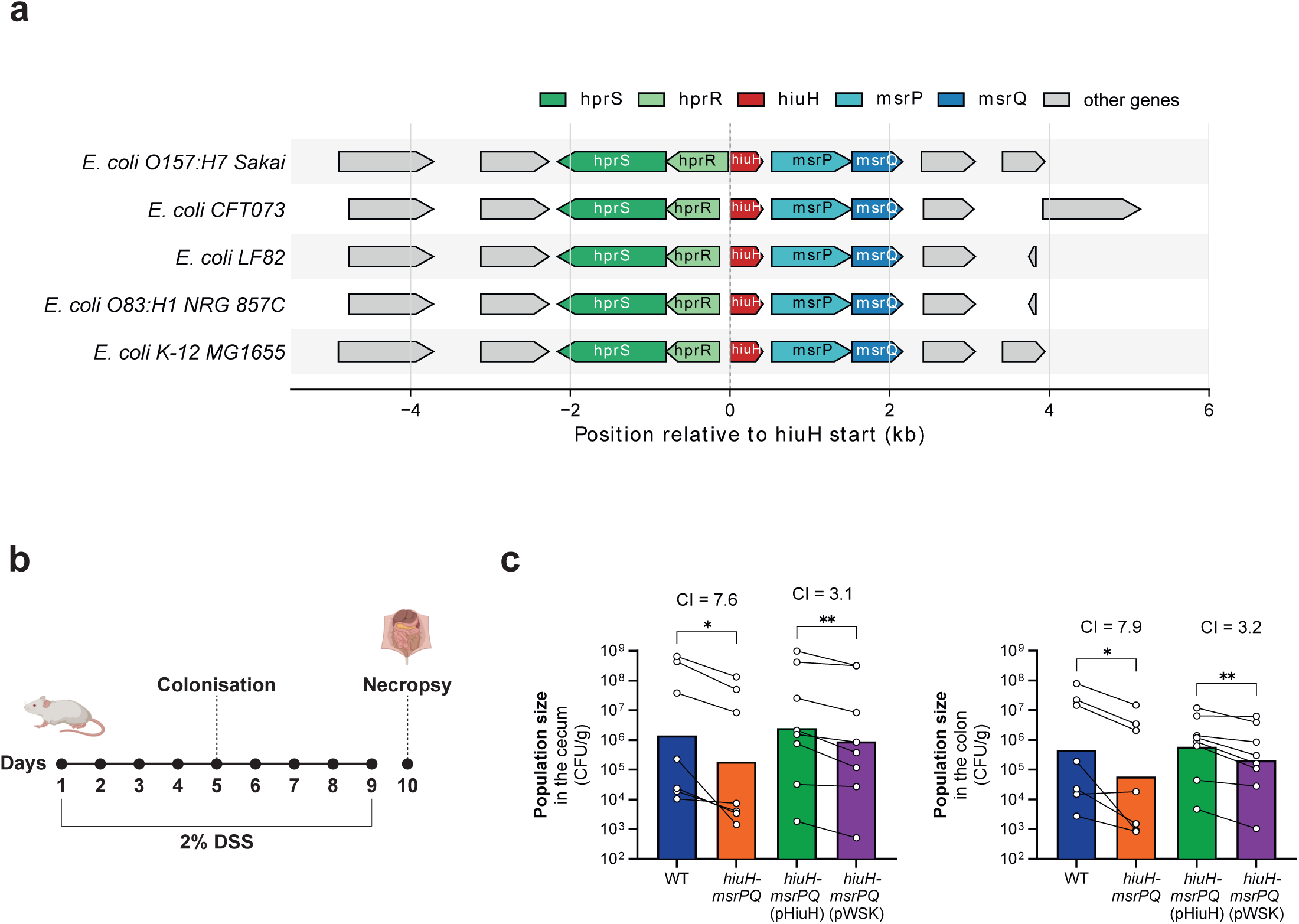
The HiuH–MsrPQ system promotes bacterial fitness in the inflamed intestine. **a.** Distribution of the *hprSR*, *hiuH* and *msrPQ* genes among various *E. coli* strains. b. Experimental design of the murine colitis model. Mice were treated with 2% dextran sodium sulfate (DSS) to induce intestinal inflammation and then experimentally colonised with *E. coli*. One group of mice received a 1:1 mixture of *E. coli* Nissle 1917 wild-type (WT) and the Δ*hiuH-msrPQ* mutant, and one group received a 1:1 of the Δ*hiuH-msrPQ* mutant harbouring pWSK129 (pWSK) and the Δ*hiuH-msrPQ* mutant harbouring pHiuH. c. Bacterial loads recovered from the cecum and colon were quantified after infection. Each dot represents the bacterial population recovered from one mouse; WT and Δ*hiuH-msrPQ* values, and the Δ*hiuH-msrPQ* (pWSK) and Δ*hiuH-msrPQ* (pHiuH) from the same animal are connected by a line. Bars indicate the geometric mean. P values were determined using by applying a two-sided Wilcoxon signed-rank test to log-transformed data.

Thus, the conserved *E. coli hiuH-msrPQ* operon defines a specialised periplasmic defence module that couples direct oxidant detoxification to repair of oxidised methionine residues. This organisation suggests that *E. coli* occupies host-associated niches in which damage repair alone is insufficient, and where direct degradation of UH provides a selective advantage.

### The HiuH-MsrPQ system increases bacterial fitness in the inflamed intestine

We reasoned that the high specificity of the HiuH-MsrPQ system to deal with UH stress may present a unique opportunity to determine whether UH influences microbial growth inside a host.

Episodes of intestinal inflammation are associated with phylum-level changes in the gut microbiome, including a bloom of commensal Proteobacteria (Pseudomonadota)^22–24^. It is conceivable that UH is formed through the reaction of inflammatory reactive oxygen species with uric acid in this environment, and that the HiuH-MsrPQ system enables *E. coli* to overcome UH stress. We therefore determined the fitness of the *ΔhiuH-msrPQ* mutant in a murine model of colitis (**Fig. 4b,c**). Since *E. coli* K-12 does not readily colonize the murine gut, we generated a *ΔhiuH-msrPQ* mutant in the human *E. coli* strain Nissle 1917 (EcN). C57BL/6 mice received dextran sulfate sodium in the drinking water. After induction of colitis, mice were colonised with an equal mixture of the EcN wild-type strain and an isogenic *ΔhiuH-msrPQ* mutant. Five days after experimental colonisation, the population size of each strain in the cecum and colon content was determined (**Fig. 4b**). Compared to the population size of the wild-type strain, the population of the *ΔhiuH-msrPQ* mutant was significantly reduced by approximately 7-fold in both the cecum and colon (**Fig. 4c**).

To determine whether this defect could be attributed to loss of HiuH function, we performed a complementation experiment in which the *ΔhiuH-msrPQ* mutant expressing HiuH was competed against the same mutant carrying an empty vector. Expression of HiuH significantly improved bacterial fitness in the inflamed gut, with a competitive index of 3.2 relative to the empty-vector control (**Fig. 4c**). Thus, these data show that HiuH promotes *E. coli* fitness during colonization of the inflamed gut.

## DISCUSSION

A central question raised by our work is to what extent UH can shape host-microbe interactions. Direct quantification of UH in vivo remains challenging because of its reactivity and transient nautre^17^, but the biological importance of an oxidant is not determined by its lifetime alone. Superoxide is highly transient, yet the widespread evolution of superoxide dismutases established that fleeting oxidants can impose strong selective pressures on cells. In this light, the conservation of the *hprS-hprR-hiuH-msrP-msrQ* architecture across *E. coli* strains suggests that UH is not merely an incidental by-product of inflammation, but a stress that bacteria have evolved to counteract. Consistent with this view, loss of the HiuH-MsrPQ module reduces *E. coli* fitness during intestinal inflammation, whereas expression of HiuH improves colonisation of the inflamed gut. Thus, the convergence of biochemical, single-cell, evolutionary and animal data supports a model in which transient urate-derived oxidative chemistry shapes bacterial fitness during inflammation.

MPO is best known for producing hypochlorous acid, but urate can compete with chloride for MPO-dependent oxidation, decreasing HOCl formation while promoting urate-derived oxidants such as UH^6^. Thus, in urate-rich inflammatory environments, the oxidative burst may be partially redirected from chlorine-based chemistry toward urate oxidation, making UH a potentially relevant extracellular oxidant at mucosal and vascular interfaces. In line with this possibility, UH has been shown to oxidise endothelial surface proteins^9^, raising the question of whether HiuH-like human enzymes could also protect from UH toxicity.

Beyond infection biology, these findings may also have implications for urate-driven human diseases. Hyperuricaemic disorders such as gout are defined by elevated urate levels, yet whether UH forms in these pathological settings remains unknown. This gap partly reflects the difficulty of measuring UH in complex samples such as synovial fluid. The rapid, specific and robust activation of the HprSR-dependent *hiuH-msrPQ* response therefore provides a unique basis for developing a bacterial biosensor to detect UH in gout and related inflammatory diseases. In parallel, the discovery of HiuH as a dedicated UH-degrading enzyme opens the possibility that UH detoxification itself could be exploited to potentially limit, urate-derived oxidative damage in disease.

## METHODS

### Strains, plasmids and primers

Bacterial strains, plasmids and primers used in this study are listed in Supplementary Tables 3, 4 and 5.

For microfluidic-based microscopy experiments, we used an *E. coli* MG1655 strain that constitutively expresses the red fluorophore mKate2, allowing cell segmentation and monitoring of cell lysis. The construct P_RNAI_-mKate2 was inserted at the chromosomal attTn7 site. To generate this strain, P_RNAI_-mKate2 amplified from strain MV6^25^ was cloned into the vector pGRG25 (addgene #16665) by XhoI/NotI restriction-ligation to create pMV65, which was then transformed into MG1655. Chromosomal insertion at the attTn7 site was performed following the protocol described in^26^.

Mutant strains derived from MV221 were generated by P1 phage transduction and selected for chloramphenicol (Cm) or kanamycin (Kn) resistance. Phage lysates were prepared from previously described donor strains: Δ*msrP*::Kn^18^, Δ*hiuH-msrPQ*::Cm^27^, and Δ*hprSR*^14^. When required, antibiotic resistance cassettes flanked by frt sites were excised by expressing Flp recombinase from plasmid pCP20.

Strains MV265 (MG1655, *ΔmsrA, ΔmsrB, ΔmsrC, ΔbisC*::Kn, P_RNAI_-mKate2::attTn7) and MV248 (MG1655 *hiuH*\*, P_RNAI_-mKate2::attTn7) were constructed by inserting P_RNAI-_mKate2 into strain BE884 (MG1655, *ΔmsrA, ΔmsrB, ΔmsrC, ΔbisC*::Kn)^18^ and LL1913 (a markerless MG1655 variant containing three stop codons immediately downstream of the *hiuH* start codon)^27^ respectively, as described above for MV221.

Plasmids pLL205, pLL198, pLL200 and pLL366 were generated by inserting yedX^6His^, yedX^Y111F-6His^, yedX^6His^ lacking the signal sequence, and pucM^6His^, respectively, into the pBAD vector (pLL21). Inserts were cloned by restriction–ligation using NdeI/XhoI (pLL205, pLL198) or EcoRI/XhoI (pLL366, pLL200) following digestion of both insert and vector with the corresponding enzymes.

Plasmids pMV26 (pUA66-P*_hiuH_*-GFP) and pMV10 (pUA66-P*_grxA_*-sfGFP) were derived from the Zaslaver promoter library^28^. Plasmid pMV41 (pUA66-P*_hiuH_*-mYPet) was derived from pMV10 and constructed by Gibson assembly to replace the GFP sequence with mYPet^29^, and the nucleotides encoding the N-terminal region of HiuH were removed to prevent periplasmic targeting of mYPet.

To generate plasmid pNT022, the region upstream of *hiuH* and the region downstream of *msrQ* were amplified from the EcN genome by PCR, and inserted into pGP706^30^ using the Gibson assembly (New England Biolabs). pNT022 was introduced into the EcN genome by conjugation, using S17-1 λ*pir* as the donor strain and by selecting for Kanamycin resistance as described in^31^. Plasmid pHiuH was generated by amplifying the hiuH gene including its native promoter region from the EcN chromosome, and inserting this DNA fragment into EcoRI-digested plasmid pWSK29^32^.

### Bacterial culture

Strains were streaked from frozen glycerol stocks onto LB agar containing the appropriate antibiotics. A single colony was used to inoculate M9 minimal medium containing M9 salts (M6030, Merck), 2 mM MgSO_4_, 0.1 mM CaCl_2_, 0.5 µg/ml thiamine, MEM amino acids (12519059, Fisher Scientific), 0.1 mg/ml L-proline, and 0.2% (w/v) glucose. Cultures were grown overnight to stationary phase at 37 °C, diluted 1:100 into fresh M9 medium, and grown to an OD_600_ of 0.2 before performing the microfluidic-based imaging experiments. For plasmid-based induction, arabinose was added to a final concentration of 0.2% (w/v) or IPTG to 100 µM.

### UH synthesis

In this study, UH was generated using riboflavin-mediated photooxidation of uric acid. Two complementary experimental configurations were employed. Batch UH synthesis was used for calibration, bolus treatments of bacterial cultures, and in vitro enzymatic assays. In parallel, a continuous-flow UH generation system was developed to enable sustained delivery of UH under microfluidic conditions compatible with long-term single-cell imaging. UH production was based on published protocols^15,17^, and spontaneous conversion of UH to HIU^17^ was carefully monitor over time.

### Batch UH synthesis

Reaction mixtures were prepared to final concentrations of 1.5 mM uric acid and 5 µM riboflavin in PBS 1X (pH 7.4 adjusted with HCl). Photooxidation reactions were carried out at 4 °C in an opaque temperature-controlled refrigerated chamber equipped with a blue-light source (Duris P5 16 Strip Deep Blue, 180 mW) centred at 455 nm. Irradiation time was adjusted according to reaction volume, with 500 µL irradiated for 5 min, 1 mL for 10 min, and 2 mL for 20 min. Reactions were performed in open weighing dishes, allowing uniform light exposure and efficient oxygen exchange to ensure sustained oxygen availability throughout irradiation. UH, HIU and residual uric acid were quantified by high-performance liquid chromatography. Separation was performed using a NucleoShell RP-18 column (250 × 4.6 mm, 5 µm) coupled to a Nucleodur C8 column (250 × 4.6 mm, 5 µm). The mobile phase consisted of solvent A (20 mM phosphate buffer, pH 7.0) and solvent B (acetonitrile), delivered at a flow rate of 0.8 mL·min^-1^. The gradient started at 100% solvent A for 15 minutes, followed by a linear transition to 30% solvent A at 23 minutes, maintained until 30 minutes, returned to 100% solvent A at 32 minutes, and equilibrated until 40 minutes. Detection was performed using a diode-array UV-visible detector. Uric acid was monitored at 290 nm, whereas UH and HIU were quantified at 308 nm. Concentrations were calculated using calibration curves generated from uric acid standards and published molar extinction coefficients of 12,300 M^-1^.cm^-1^ at 290 nm for uric acid and 6,540 M^-1^.cm^-1^ at 308 nm for UH and HIU.

Reaction mixtures were further analysed by mass spectrometry before and after irradiation to confirm UH formation. Analyses were performed using a SYNAPT G2 HDMS mass spectrometer (Waters) equipped with an atmospheric pressure ionisation source. Samples were ionised in negative electrospray mode with a spray voltage of −2.27 kV and a cone voltage of −20 V. Nitrogen was used as the nebulising gas at a flow rate of 100 L·h^-1^ and mass spectra were acquired using a time-of-flight analyser. Prior to analysis, samples were diluted tenfold in methanol and introduced into the ion source by direct infusion at 10 µL·min^-1^.

### Continuous UH synthesis

To enable sustained UH exposure, a continuous-flow UH synthesis system was implemented. A solution consisting of M9 minimal medium supplemented with glucose and 5 µM riboflavin and 500 µM uric acid was pumped at a flow rate of 2 mL.h^-1^ through opaque tubing measuring 100 cm in length with an inner diameter of 0.02 inches. The tubing passed through a light-tight enclosure equipped with a switchable 15 cm LED strip emitting at 455 nm (Duris P5 16 Strip Deep Blue). We calculated the residence time within the illuminated region ∼55 s. The UH-containing medium was then delivered directly to a microfluidic device for live-cell imaging at 100× magnification. The entire setup was operated in a temperature-controlled dark room maintained at 22 °C. UH, HIU and uric acid concentrations were measured by HPLC at the outlet of the tubing, immediately upstream of the microfluidic chip. Measurements were performed in triplicate across at least 3 independent experiments. The resulting UH concentrations were stable over time within experimental error and are summarised in Extended Data Fig. 2.

### Beta-galactosidase assay

Cultures of strain LL1700^27^ were grown in M9 medium supplemented with glucose and adjusted to an OD_600_ of 0.3. UH, preincubated or not with 0.50 µM HiuH or HiuH^Y111F^, was added to a final concentration of 86 µM, and cultures were incubated for 1 h at 37 °C. β-galactosidase activity was measured as previously described^27^.

### Immunoblotting

Cells were lysed in Laemmli buffer (2% SDS, 10% glycerol, 60 mM Tris-HCl pH 7.4, 0.01% bromophenol blue, 50mM DTT and 5% β-mercaptoethanol), and total protein amounts were normalised to the culture optical density at 600 nm. Samples were heated for 10 min at 95 °C and resolved by SDS-PAGE. Proteins were transferred onto PVDF or 0.2 µm nitrocellulose membranes and probed with guinea pig anti-MsrP, rabbit anti-HiuH, or rabbit anti-RecA primary antibodies, followed by HRP-conjugated anti–guinea pig or anti–rabbit IgG secondary antibodies (Promega). Signals were detected by chemiluminescence using an ImageQuant LAS4000 system (GE Healthcare Life Sciences). Uncropped immunoblots are provided in Supplementary Fig.2.

### Fractionation

Sphaeroplasting was performed using the *hrpS^M153A^* strain, which exhibits constitutive expression of the *hiuH–msrPQ* operon^14^. A 2 ml culture sample was centrifuged, washed, and resuspended in 250 µl of 0.2 M Tris-HCl buffer (pH 8). The suspension was mixed with 250 µl of 0.2 M Tris-HCl (pH 8) containing 1 M sucrose and 1 mM EDTA. Lysozyme (2 µl, 15 mg/ml) was added, and the mixture was incubated at room temperature for several minutes. The suspension was then centrifuged, and the supernatant was collected as the periplasmic fraction.

### Protein oxidation

Calmodulin (6 µM), PucM (1.2 µM) or HiuH (1.2 µM) were incubated with UH (200 µM) for 15 min at room temperature in phosphate buffer. For CaM oxidation assays, reactions were performed either in the absence or presence of purified HiuH or HiuH^Y111F^ (50 µM), followed by quenching with Laemmli SDS sample buffer, heating at 95 °C for 10 min, separation on 4-20% gradient SDS–PAGE gels, and visualisation by Coomassie blue staining (for Calmodulin) or immunodetection (for PucM and HiuH).

### Protein purification

Recombinant HiuH^6His^, HiuH^Y111F-6His^, and PucM^6His^ were produced in *E. coli* MG1655 transformed with the corresponding pBAD expression plasmids. Cultures were grown in LB medium supplemented with kanamycin (100 µg/ml) at 37 °C until mid-log phase, and protein expression was induced with 0.2% (w/v) arabinose for 4 h at 37 °C. Cells were harvested by centrifugation (6,000 rpm, 5 min, 4 °C) and stored at −80 °C until use. For lysis, cell pellets were resuspended in 100 mM Tris-HCl, 50 mM NaCl (pH 8.0) and supplemented with lysozyme (0.5 mg/ml), DNase I (10 µg/ml), and RNase A (10 µg/ml). The suspension was incubated for 10 min at room temperature and disrupted using a French press. Cell debris was removed by centrifugation (8,500 rpm, 10 min, 4 °C), and the supernatant was filtered (0.45 µm) prior to affinity purification. Proteins were purified on a HisTrap Ni^2+^-NTA column (Cytiva) pre-equilibrated with buffer A (100 mM Tris-HCl, 50 mM NaCl, pH 8.0). Bound proteins were eluted using a linear gradient of buffer B (100 mM Tris-HCl, 50 mM NaCl, 1 M imidazole, pH 8.0) at a flow rate of 0.5 ml/min. Fractions containing the target protein were identified by SDS-PAGE, pooled, and concentrated using centrifugal filters (Amicon Ultra, 10 kDa cutoff). Imidazole was removed by washing three times with buffer A. Protein concentration was determined spectrophotometrically at 280 nm using theoretical extinction coefficients calculated from the amino acid sequence. Purified proteins were flash-frozen in liquid nitrogen and stored at −80 °C until use.

### Enzymatic assays

The degradation of UH and HIU was monitored spectrophotometrically by following absorbance changes at 312 nm at 30°C. Measurements were acquired every 1.2 s for 10 min using a quartz cuvette at room temperature. Immediately after thawing on ice, purified proteins (HiuH, HiuH^Y111F^, and PucM) were diluted into 50 mM potassium phosphate buffer (pH 7.4). Final protein concentrations were either 0.06 µM (1X) or 0.6 µM (10X). For UH degradation assays, UH was thawed from −80 °C stocks and immediately added to the cuvette at a final concentration of 50 µM. The solution was rapidly mixed, and data acquisition was started without delay. For HIU degradation assays, 0.06 uM of uricase from *Bacillus fastidiosus* (Sigma-Aldrich, 94310) was added to the phosphate buffer prior to protein addition, and absorbance was monitored for 1 min. Uric acid (final concentration 50 µM) was then added to initiate HIU production, and the reaction was followed for an additional 90 s. At 2 min 30 s, purified proteins were introduced at the indicated concentrations, and the degradation of HIU was monitored in real time.

### Microscopy

Time-lapse imaging was performed using a Nikon Eclipse TiE inverted fluorescence microscope equipped with a 100× NA 1.40 oil-immersion objective, motorised stage, sCMOS camera (Hamamatsu ORCA-Fusion BT), LED excitation source (Lumencor SpectraX), and a perfect focus system. The SpectraX system comprised independently controllable LED channels and a motorised filter turret equipped with monochromatic filter cubes for GFP, CFP, YFP, and mCherry. The microscope was controlled using NIS. Exposure times were 50 ms for P*_rna1_*-mKate2 (555 nm, 10% LED intensity), 50 ms for P*_hiuH_*-mYPet (510 nm, 50% LED intensity), 50 ms for msfGFP-glpT (470 nm, 50% LED intensity), 50 ms for lpoB-mCherry (555 nm, 50% LED intensity), and 50 ms for msfTq2ox (440 nm, 50% LED intensity). The microscope chamber (Okolabs) was maintained at 37 °C throughout the experiments. Images were acquired every 10 minutes, except for the dataset used to train the machine learning model, for which images were acquired every 3 minutes to increase the temporal resolution of response dynamics.

### Microfluidics

The microfluidic imaging device was based on the original mother machine design^33^. Trenches were fabricated with dimensions of 1.2 µm (width) × 1.2 µm (height) × 25 µm (length), while main flow channels measured 100 µm (width) × 25 µm (height). Each device contained three independent lanes with separate inlets and outlets, allowing simultaneous analysis of multiple experimental conditions or bacterial strains. Structures were fabricated onto silicon wafers by ConScience AB. Polydimethylsiloxane (PDMS) chips were produced using Sylgard 184 (Farnell, 101697) by mixing pre-polymer and curing agent at a 10:1 ratio in a 50 ml Falcon tube. The mixture was poured into a square Petri dish and degassed under vacuum to remove air bubbles. Subsequently, the PDMS was poured onto the wafer, previously wrapped in aluminium foil, and degassing was repeated if necessary until all bubbles were eliminated. The PDMS was then cured at 65°C for 2 h. Media inflow and outflow holes were created using a 0.75 mm biopsy punch (Fisher Scientific, 11683284). PDMS chips were cleaned by repeated taping and untaping to remove residual dust, then bonded to microscope coverslips (No. 1.5) using corona treatment (BD-A corona treater, ETP). Bonding was enhanced by placing the assembled chip in an oven at 95°C for 3 h immediately after corona exposure. Cells were loaded into the device by pipetting, followed by centrifugation at 5000 rpm for 10 min using a benchtop centrifuge to position cells within the growth channels. To minimise cell aggregation, supplemented M9 minimal medium containing 0.85 mg.ml^-1^ surfactant Pluronic F127 (P2443-250G, Sigma Aldrich) was continuously supplied through Tygon ND 100-80 microbore silicone tubing (15670211, Fisher Scientific) from 50 ml or 30 ml syringes controlled by motorised pumps (NE-300 InfusionONE, Darwin Microfluidics). After cell loading, medium was delivered at a flow rate of 2 ml.h^-1^. Cells were incubated within trenches for approximately 1 h to reach steady-state conditions prior to imaging and data acquisition.

### Imaging data analysis

Data from time-lapse imaging of mother machine experiments were analysed using CSV files exported by the BACMMAN^34^ plugin in FIJI^35^. Image pre-processing was performed in BACMMAN using the PRNAI–mKate2 fluorescence channel to register individual growth trenches, correct lateral drift in x–y coordinates, and compensate for image rotation. Trenches and cells were segmented on the basis of constitutive mKate2 fluorescence, and cell lineages were reconstructed over time using the same signal. Segmentation and tracking outputs were systematically inspected, and errors in cell boundary detection or lineage assignment were manually corrected. The resulting datasets were subsequently analysed using custom Python scripts to compute cellular parameters, as described below.

### Growth rate

The single-cell growth rate was defined and calculated by BACMMAN according to an exponential growth model. Cell size *S(t)* at time *t* was described as: *S(t)=S0.e^μt^* where *S0* corresponds to the cell size at birth, *μ* is the growth rate, and *t* is the time elapsed since birth. For each individual cell, BACMMAN estimate *μ* by fitting this model to the measured cell size trajectory over time.

### Survival and physiological state analysis

The analysis was restricted to mother cells, which were individually tracked across consecutive frames using unique identifiers. Survival was quantified as the fraction of initially present mother cells that remained detectable at each time point. Because mother cells are retained at the closed end of the trench, loss from the field of view is not expected to result from physical escape. However, survival alone does not discriminate between distinct physiological outcomes. Therefore, each surviving cell was further classified based on its growth dynamics.

Physiological states were assigned from the temporal evolution of cell length, measured using BACMMAN. Cells were classified as lysed when no length value was available at the defined time point. Filamentation was defined as a more than twofold increase in average cell length between the first and last 20% of time points following stress. Cells that neither divided nor filamented were classified as growth-arrested.

### Cell fluorescence

mYPet, sfGFP, mCherry and msfTqox fluorescence signals were quantified using the *ObjectFeatures-Mean* measurement module implemented in BACMMAN, which was edited to compute the average fluorescence intensity within individual segmented bacterial cells. The temporal delay between cytoplasmic and periplasmic release events was determined manually.

### Machine learning

We used a random forest regression model to investigate the causes of heterogeneous stress response magnitude among individual *E. coli* cells exposed to identical concentration of UH stress. Specifically, we focused on mother cells located at the closed end of each growth channel that can be observed for many generations, tracking their time-lapse trajectories across three experimental replicates. The microscopy time-lapse data was fed into BACMMAN^34^ to extract time series data as CSV files which were processed using custom Python code to create input for the machine learning algorithm. The final dataset included individual time traces of 1464 mother cells across all replicates. Next, 126 features were extracted from each of these traces for treated (63 features) and untreated (63 features) condition, as described in Choudhary et al.^21^. These features encompass 7 mathematically derived features (including kurtosis, skewness, median, mean, minimum, maximum, range) from cellular properties (including cell length, area, instantaneous elongation rate, averaged size increase rate) and features describing the environmental descriptors i.e. barrier cells (including barrier volume, surface area, surface area to volume ratio, volumetric surface area to volume ratio, total number of barrier cells). Each feature was associated with the peak magnitude of stress response after UH exposure. The dataset was then shuffled, and 80% was used to train a random forest regressor comprising 100 tress and a depth of 2000. The remaining 20% (294 mother cells, unseen by the model) served as the test set. We evaluated the model performance by computing accuracy as 1 - mean absolute percentage error, calculated between predicted and experimentally measured response magnitude for each mother cell. The resulting model achieved an accuracy of 62.53%. We identified which features contributed the most to model’s predictive ability by applying the mean decrease impurity test using the scikit-learn library in python.

### Bioinformatics

To examine HiuH proteins across bacteria, we first extracted a large-scale dataset of 15,389 sequences containing Transthyretin/hydroxyisourate hydrolase domain from the InterPro database (IPR023416)^36^. Then, we identified signal peptide sequences, sub-cellular localisations and methionine statistics of each protein sequence using singlaIP^37^, CELLO^38,39^ and EMBOSS packages, respectively. We then carried out an all-vs-all sequence similarity searching and clustering (identity ≥ 50 % and bidirectional coverage ≥ 70 %) of the dataset using MMseqs2 algorithm^40,41^. A total of 17 largest clusters gathering in 12,540 sequences (81%) were retained, and sequences of each cluster were then subjected to multiple sequence alignments (MSA) and examined using JalView software. These data made possible the selection of a last dataset of 54 distant HiuH proteins of 39 relevant bacterial strains (Table S1), which may represent members of the 17 largest clusters, using several criteria: non-redundant and divergent sequences, either from unique or duplicated genes, variabilities of protein features and diversity of bacterial lineages. Both uniqueness and duplication of the *hiuH* genes were also checked by searching homologues of the HiuH reference sequence of *E. coli* K-12 (UniProtKB acc. P76341) against RefSeq genomic database from NCBI (https://www.ncbi.nlm.nih.gov/) using both BlastP and TblastN. Our dataset was selected in order to establish possible evolutionary relationships between phylogeny, sequence features and genomic contexts of HiuH proteins across diverse lineages. The phylogenetic tree was computed using ModelTest-NG, Maximum Likelihood method (ML), LG+G model under raxmlGUI software version 2.0^42^ after performing a MSA of HiuH proteins using JalView tool and TrimAl algorithm^43^. The nodes robustness was estimated through Bootstrap (BP) analyses of 1,000 replicates. The comparative analysis of hiuH genomic contexts in complete genomes was carried out using FlaGs2 software^44^, which used HiuH proteins and RefSeq genome assembly as input datasets and clustered a set of 432 hiuH neighbourhood genes (4 upstream and 4 downstream) into homologous groups using sensitive Hidden Markov Model-based method Jackhmmer (E-value cut-off = 1e^-10^).

### Murine colitis model

All experiments were conducted in accordance with the policies of the Institutional Animal Care and Use Committee at UC Davis. C57BL/6J mice were bred at UC Davis under specific pathogen-free conditions. All mice were on a 12h light/dark cycle and consumed food (Teklad 2918) and sterile water (Hydropac) ad libitum. Male mice, 8-9 weeks of age, received sterile-filtered dextran sulfate sodium (DSS; 2% w/v) in the drinking water in a clean water bottle for four days. To colonise mice, *E. coli* Nissle 1917 wild-type and the isogenic Δ*hiuH-msrPQ* mutant, harboring plasmids pWSK29 and pWSK129 respectively, were used. Similarly, cultures of the Δ*hiuH-msrPQ* mutant, either harboring pWSK129 or pHiuH were used. Bacterial strains were grown overnight (16h) in 100 mL of LB broth. Cultures were centrifuged and resuspended in fresh LB to create a suspension of 10^10^ CFU/mL. Cultures were then mixed in 1:1 ratio. Mice received 0.1 ml of this suspension, corresponding to 5 x 10^8^ CFU for each strain, by oral gavage. The inoculum was plated on selective agar plates. DSS treatment was continued for four days. Mice then received fresh drinking water for another day. After mice were euthanized, content from the colon and cecum was collected in cold phosphate buffered saline. Suspensions were plated on selective agar plates. The competitive index (CI), a measure of fitness, was calculated by dividing the population size of the wild-type strain by the population size of the mutant in each organ for each animal. This ratio was then corrected by the input ratio of wild-type and mutant bacteria.

## DATA AVAILABILITY

Data supporting the findings of this study will be made available upon publication. Source data will be provided with the paper.

## AKNOWLEDGEMENTS

We thank members of the Ezraty lab for insightful discussions. We thank Bastien Charrat for his help with the construction of strain MV221. We thank Jean-François Collet and Michael Deghelt for pMV53 and pMV54 plasmids gift. This work was supported by: Agence Nationale de la Recherche, ANR JCJC AMORE (ANR-24-CE44-2908) and IM2B (AO-IM2B-NE-2024-02-VINCENT) to M.S.V, and ANR NeutrOX (ANR-21-CE15-0039) to B.E. Work in S.E.W.’s lab is supported by the NIH (AI188307, AI118807, AI171537); the funders had no role in study design, data collection and interpretation, or the decision to submit the work for publication. Any opinions, findings, conclusions, or recommendations expressed in this material are those of the authors and do not necessarily reflect the views of the funding agencies.

## AUTHOR INFORMATION

**M.S.V.** performed strain construction and microfluidic-based experiments, analysed the data, supervised the study, acquired funding, contributed to conceptualisation, generated figures, and wrote the manuscript. **L.L.** performed strain construction and biochemical experiments, including β-galactosidase assays, protein purification, protein oxidation, and enzymatic activity assays, and contributed to data analysis. **M.J.** collected datasets for machine learning and performed data analysis. **P.S.** conducted HPLC, mass spectrometry, and chemical assay analyses. **KEK**. Carried out bioinformatic analyses. **M.C.** carried out in vitro experiments, including protein purification, protein oxidation and enzymatic activity assays. **D.C.** developed the machine learning model and contributed to data analysis. **N.V.** and **S.P.** provided intellectual input on chemical analyses. **M.G.W, N.T** and **S.E.W** conducted and analysed mice infection experiments. **B.E.** initiated the investigation on urate hydroperoxide, contributed to data analysis and conceptualisation, supervised the study, and acquired funding.

---

**Extended Data Figure 1:**
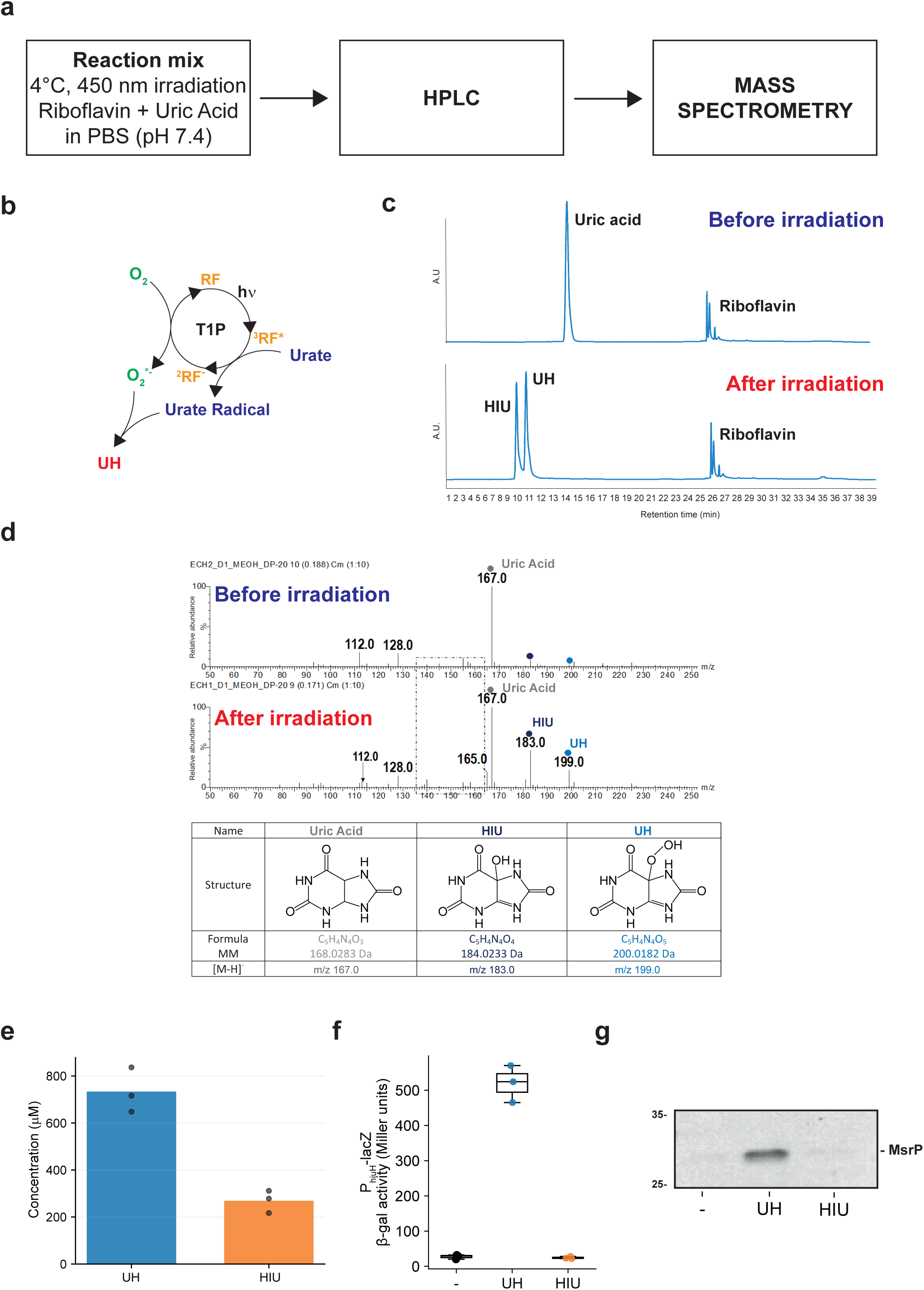
Batch production and quantification of UH. **a.** Schematic of the pipeline used for the batch production and quantification of UH. b. Proposed reaction mechanism for UH formation via Type 1 photooxidation (T1P) mediated by riboflavin (RF), based on references^15–17^. c. High-performance liquid chromatography (HPLC) chromatograms of the reaction mixture before (top) and after (bottom) irradiation. New peaks corresponding to UH and HIU appear post-irradiation. d. Mass spectrometry spectra of the reaction mixture before (top) and after (bottom) irradiation. Spectra reveal the presence of uric acid (m/z 167.0) and the formation of two new species, HIU (m/z 183.0) and UH (m/z 199.0), after irradiation. The table summarizes the molecular structures, exact masses, and detected m/z values of uric acid, HIU, and UH. e. Quantification of UH and HIU concentrations by calculating the area under the HPLC curve. Three experimental replicates are shown, indicating that UH is the predominant species post-irradiation. The UH/HIU concentration ratios across replicates are 0.72, 0.72, and 0.75. f. UH and HIU peaks were separated by HPLC, isolated, and collected individually. The resulting purified fractions were used to measure β-galactosidase activity from a chromosomally integrated P_hiuH_-*lacZ* transcriptional fusion (ectopic locus) in cells exposed to no treatment (–), 10 μM UH, or 10 μM HIU. Box plots show data from three independent biological replicates; individual data points are overlaid. g. Cells used in panel f were analysed by immunoblotting using anti-MsrP antibodies. Molecular weight markers (kDa) are indicated

**Extended Data Figure 2:**
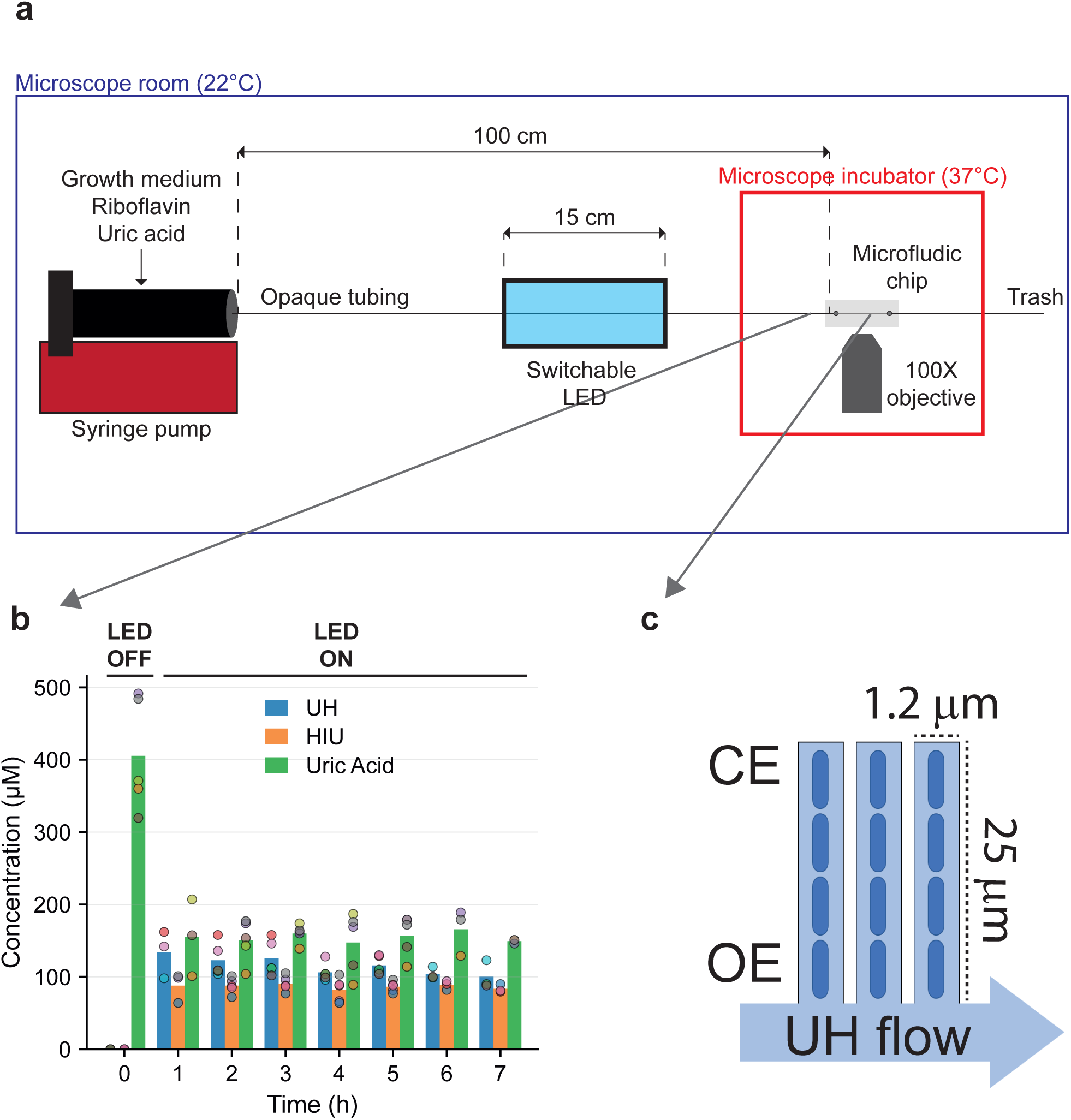
Microfluidic-based setup for continuous-flow synthesis and delivery of UH. **a.** Schematic of the continuous UH synthesis system coupled to live-cell imaging (not to scale). b. UH, HIU and uric acid concentrations at the tubing outlet were quantified by HPLC. The plot shows mean ± standard deviation from at least three independent replicates. LED was turned off at time 0. c. Design of the mother machine-type microfluidic chip used in this study. Cells grow inside channels that are 25 µm long and 1.2 µm wide. CE, closed end; OE, open end.

**Extended Data Figure 3:**
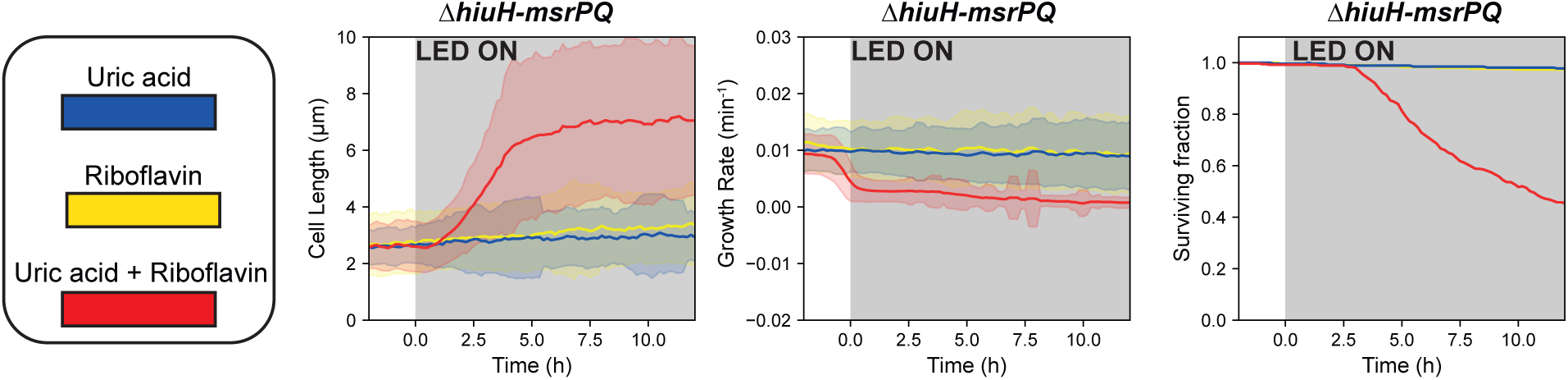
Irradiation of riboflavin or uric acid alone does not affect bacterial physiology. Cell length, growth rate, and survival of the *ΔhiuH-msrPQ* strain are shown under constant 455 nm LED exposure (100 mA; LED ON: shaded area) with uric acid only (blue), riboflavin only (yellow), or both (red).

**Extended Data Figure 4:**
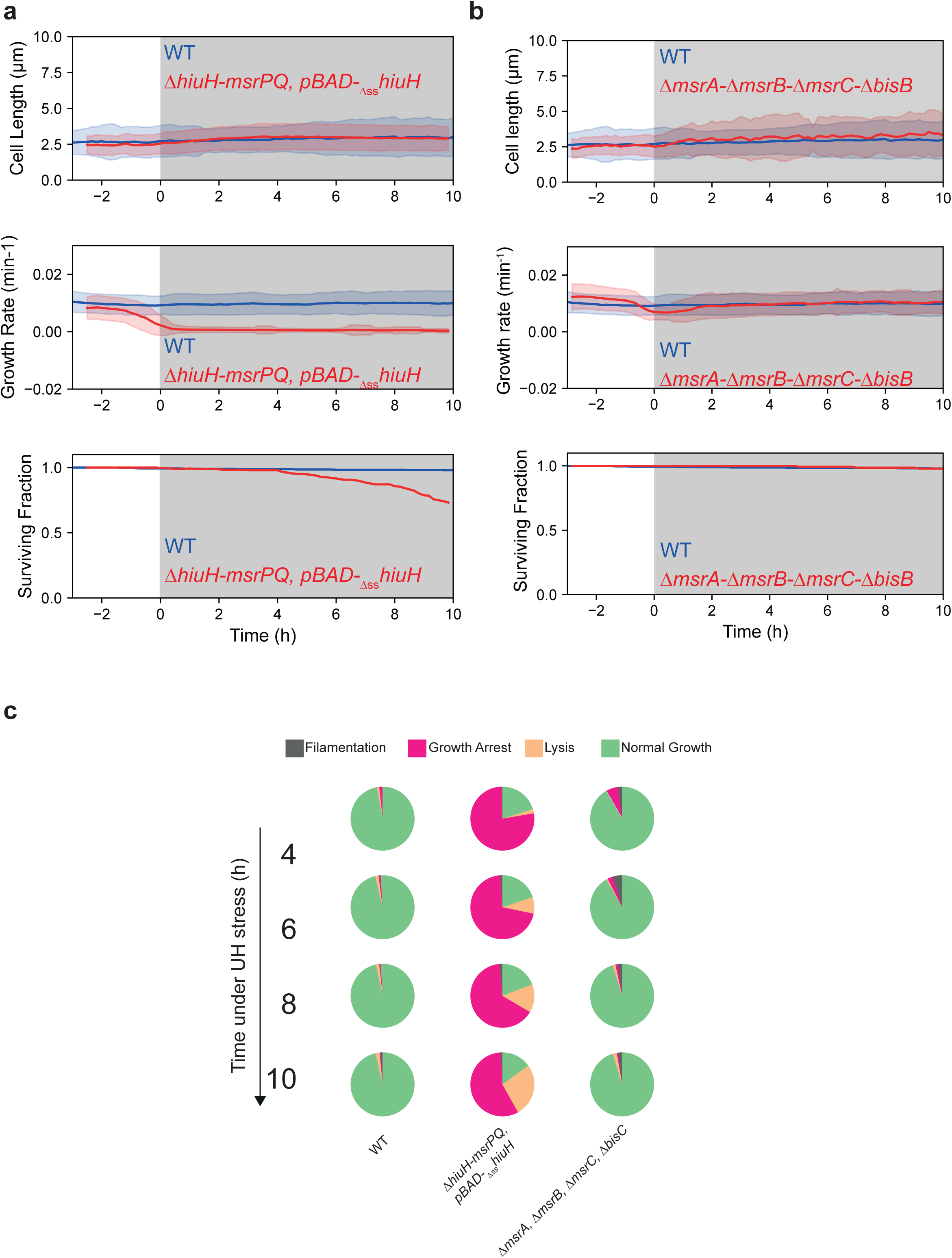
HiuH periplasmic localisation mediates UH tolerance. **a,b,** Population-averaged cell length (top), growth rate (middle), and survival (bottom) over time in WT and *ΔhiuH-msrP* expressing cytoplasmic HiuH (pBAD-*_Δss_hiuH*) (a) and in WT (n=585) and Δ*msrA ΔmsrB ΔmsrC ΔbisC* cells (n=145) (b) under constant UH treatment (shaded area). c. Quantification of physiological state fractions over time for the same strains and conditions shown in panel a and b.

**Extended Data Figure 5:**
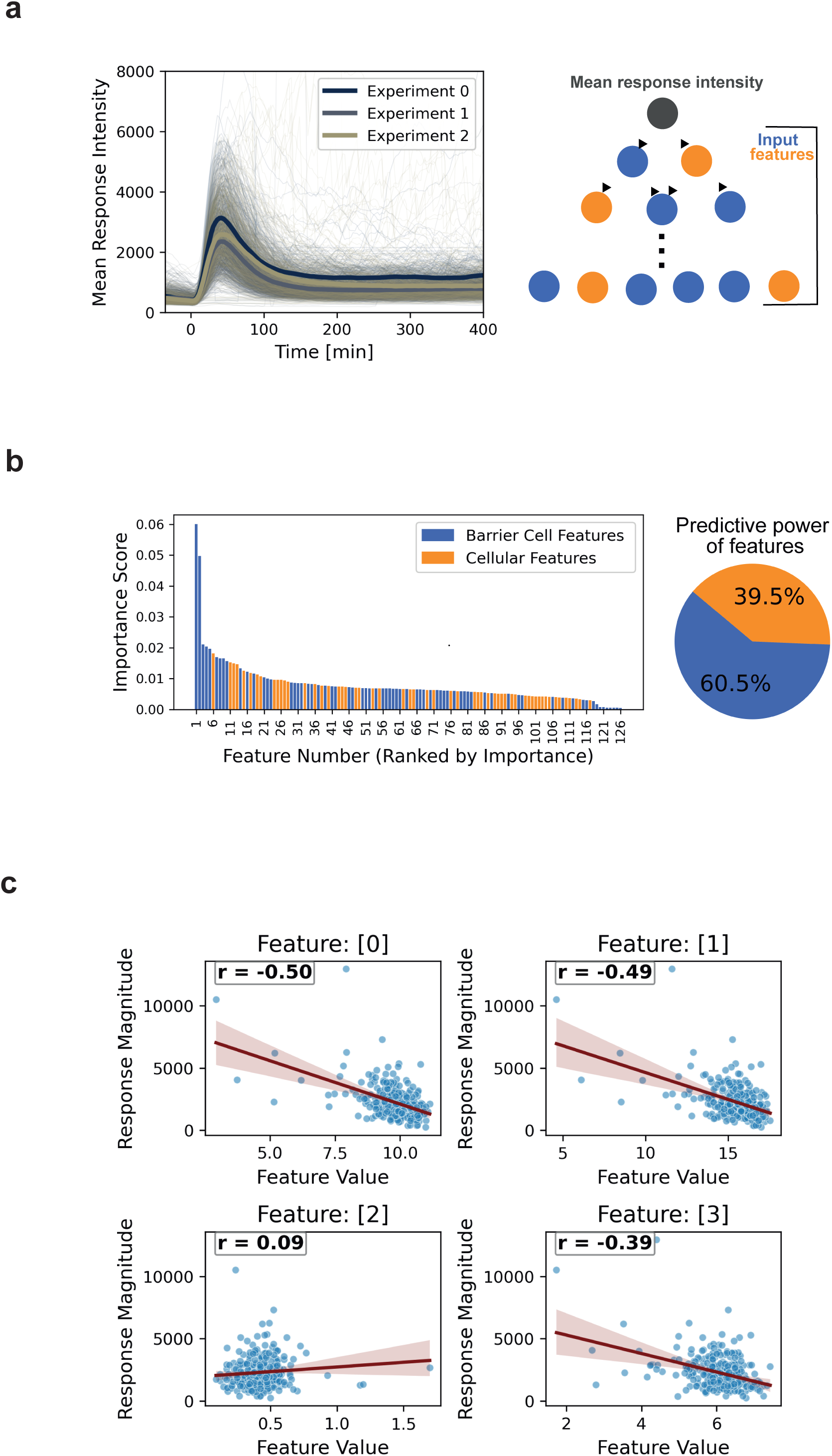
Collective cell-cell protection drives UH stress response heterogeneity. **a.** (left) Mean response intensity to UH treatment over time from t = 0 min. Bold lines represent response of mother cells for three independent replicates, while thin lines show individual single cell traces. (Right) Schematic of a random forest architecture illustrating the prediction of mean response intensity based on input features from the focal cell (orange) and its surrounding barrier cells (blue) b. (left) Feature importance ranking depicting the relative contribution of various features to the model’s predictive power. Features derived from barrier cells are shown in blue and focal cell features are marked in orange. (right) Pie chart summarizing the aggregate importance of barrier cell versus focal cell features in the machine learning model. c. The scatter plots represent the relationship between the response magnitude and the values for the first four most important features identified by the model. R values denote Pearson correlation coefficient in each plot and the red line denotes the linear regression relating the feature value and the response magnitude.

**Extended Data Figure 6:**
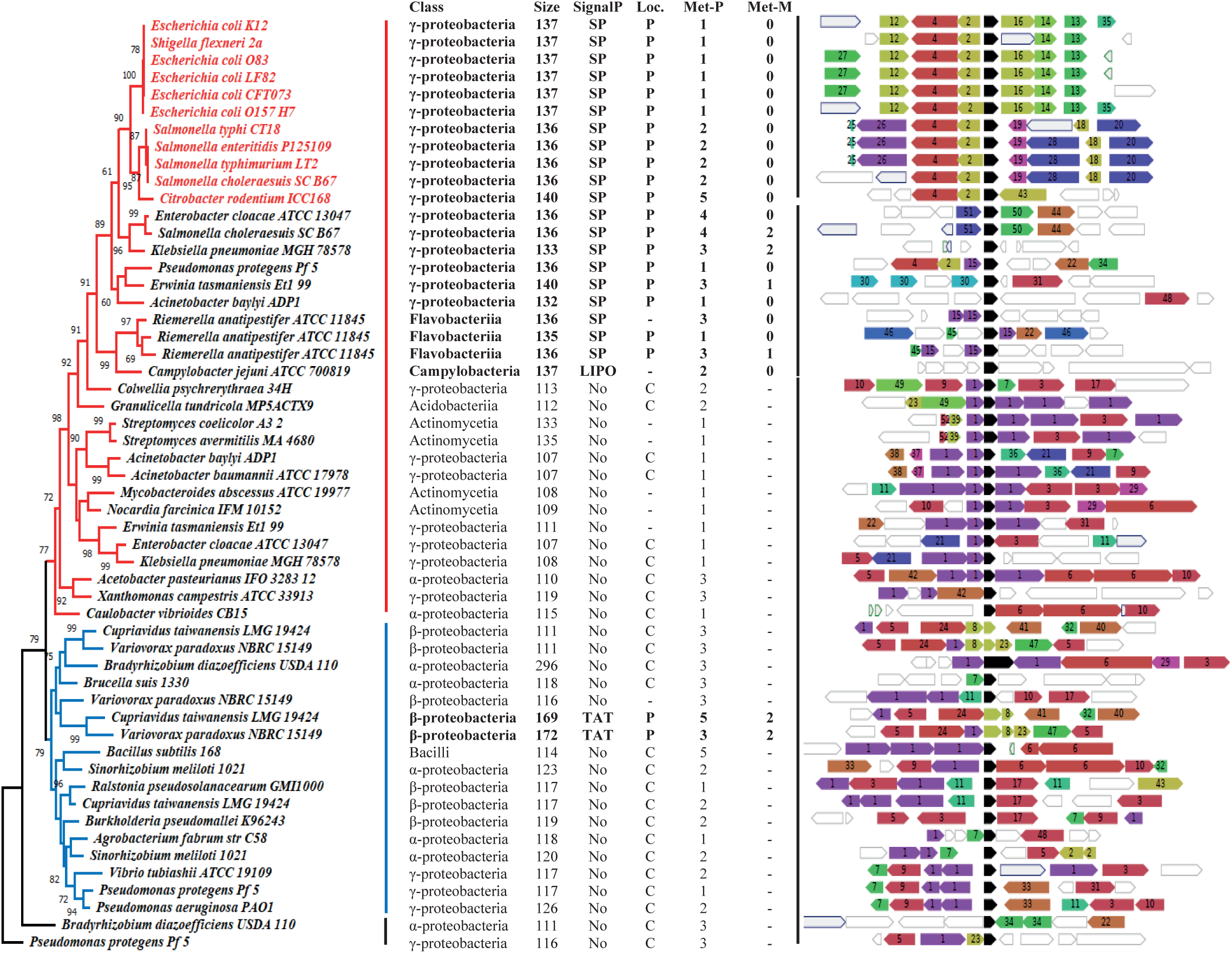
Phylogenetic tree, sequence features, and genomic contexts of HiuH proteins in diverse bacterial lineages. On the left, the phylogenetic tree shows bootstrap support values obtained from 1,000 replicates, reported next to the branch nodes. In the middle, sequence features identified in HiuH proteins are shown: SignalP prediction, indicating the type of signal peptide detected: SP, LIPO, or TAT; Loc. indicating subcellular localisation: periplasmic, cytoplasmic, or unknown (-); Met-P indicating the number of methionine residues detected in the HiuH primary sequence; and Met-M indicating the number of methionine residues detected in the mature HiuH protein. On the right, genomic contexts of *hiuH* genes across bacteria are shown. Identical numbers and colours represent clusters of potential homologous genes flanking *hiuH* genes, whereas genes without numbers or colours lack homologues among the flanking genes analysed. Black denotes the *hiuH* gene, assigned to cluster number 53.

**Table S1.** List of bacterial strains and accession numbers for their genome assemblies and HiuH proteins from RefSeq, Swissprot and UniProt databases

**Table S2.** Flanking genes associated with HiuH query genes retrieved from the RefSeq database using FlaGs

**Table S3:** Strains used in this study.

| Laboratory reference | Genetic background and relevant genotype | Plasmid | Source |
| --- | --- | --- | --- |
| MV8 | Escherichia coli K12 MG1655 |  | Laboratory collection |
| LL1700 | MG1655 lacI-yedW::cat P<sub>hiuH</sub>::lacZ |  | Loiseau et al., Mol Mic 2022 |
| MV6 | Escherichia coli K12 AB1157, flhD-, P<sub>RNAI</sub>-mKate2::attTn7, alkA-mYPet, ada-CFP::kn |  | Vincent et Uphoff., NAR 2021 |
| BE870 | MG1655 | pBBR- hprS<sup>M153A</sup> | El Hajj et al., J Bac 2022 |
| LL1913 | MG1655, *hiuH* * |  | Loiseau et al., Mol Mic 2022 |
| BE996 | MG1655, Δ*hiuH*-Δ*msrPQ* | | Loiseau et al., Mol Mic 2022 |
| MV221 | MG1655, P<sub>RNAI</sub>-mKate2::attTn7 |  | This study |
| MV248 | MG1655, *hiuH* *, P<sub>RNAI</sub> -mKate2::attTn7 |  | This study |
| MV257 | MG1655, *hiuH* *, P<sub>RNAI</sub> -mKate2::attTn7 | pBAD-*hiuH*<sup>6His</sup> | This study |
| MV242 | MG1655, Δ*hiuH*-Δ*msrPQ*::CmR, P<sub>RNAI</sub>-mKate2::attTn7 | | This study |
| BE1955 | MG1655, Δ*hiuH*-Δ*msrPQ*::CmR, P<sub>RNAI</sub>-mKate2::attTn7 | pBAD | This study |
| MV245 | MG1655, Δ*hiuH*-Δ*msrPQ*::CmR, P<sub>RNAI</sub>-mKate2::attTn7 | pBAD-*hiuH*<sup>6His</sup> | This study |
| BE1957 | MG1655, Δ*hiuH*-Δ*msrPQ*::CmR, P<sub>RNAI</sub>-mKate2::attTn7 | pBAD-Δ*sshiuH*<sup>6His</sup> | This study |
| MV256 | MG1655, Δ*hiuH*-Δ*msrPQ*::CmR, P<sub>RNAI</sub>-mKate2::attTn7 | pBAD-pucM<sup>6His</sup> | This study |
| MV238 | MG1655, Δ*msrP*, P<sub>RNAI</sub>-mKate2::attTn7 | | This study |
| MV266 | MG1655, ΔHprSR, P<sub>RNAI</sub>-mKate2::attTn7 | | This study |
| MV258 | MG1655 | pTHV037-SP<sub>dsbA</sub>TqOX-CytomCherry | This study |
| MV261 | MG1655, ΔhiuH-ΔmsrPQ | pTHV037-SP<sub>dsbA</sub>TqOX-CytomCherry | This study |
| MV204 | MG1655 | pTHV037-lpoB-mcherry-msfGFP-glpT | This study |
| MV262 | MG1655, ΔhiuH-ΔmsrPQ | pTHV037-lpoB-mcherry-msfGFP-glpT | This study |
| MV265 | MG1655, ΔmsrA, ΔmsrB, ΔmsrC, ΔbisC::KanR, P<sub>RNAI</sub>-mKate2::attTn7 | | This study |
| MV230 | MG1655, P<sub>RNAI</sub>-mKate2::attTn7 | pMV41 (pUA66-P<sub>hiuH</sub>-mYPet) | This study |
| MV251 | MG1655, hiuH*, P<sub>RNAI</sub>-mKate2::attTn7 | pMV41 (pUA66-P<sub>hiuH</sub>-mYPet) | This study |
| MV264 | MG1655, ΔmsrP, P<sub>RNAI</sub>-mKate2::attTn7 | pMV41 (pUA66-P<sub>hiuH</sub>-mYPet) | This study |
| MV240 | MG1655, P<sub>RNAI</sub>-mKate2::attTn7 | pMV26 (pUA66-P<sub>grxA</sub>-SfGFP) | This study |
| EcN | *E. coli* Nissle 1917 wild-type strain |  | Grozdanov et al., J Bact, 2004 |
| NT98 | EcN ΔhiuH-msrPQ | | This study |

**Table S4:**
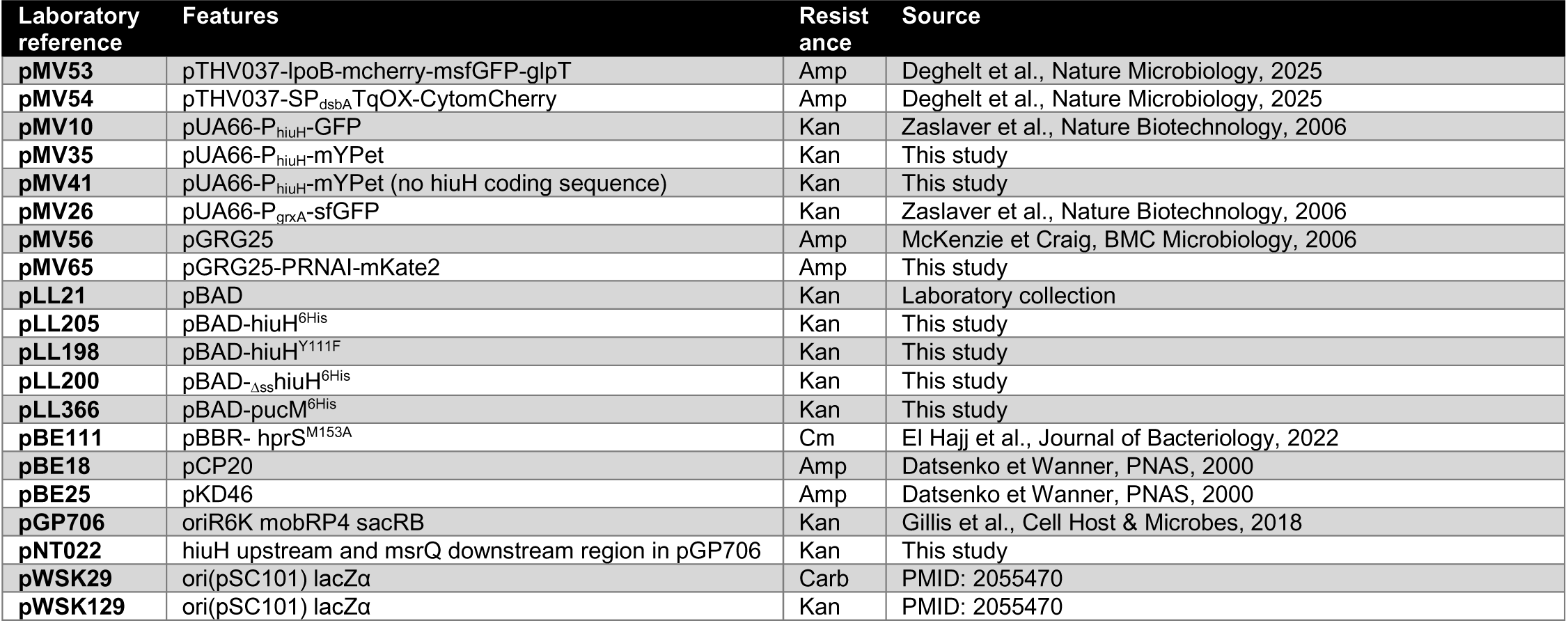
Plasmids used in this study.

**Table S5:**
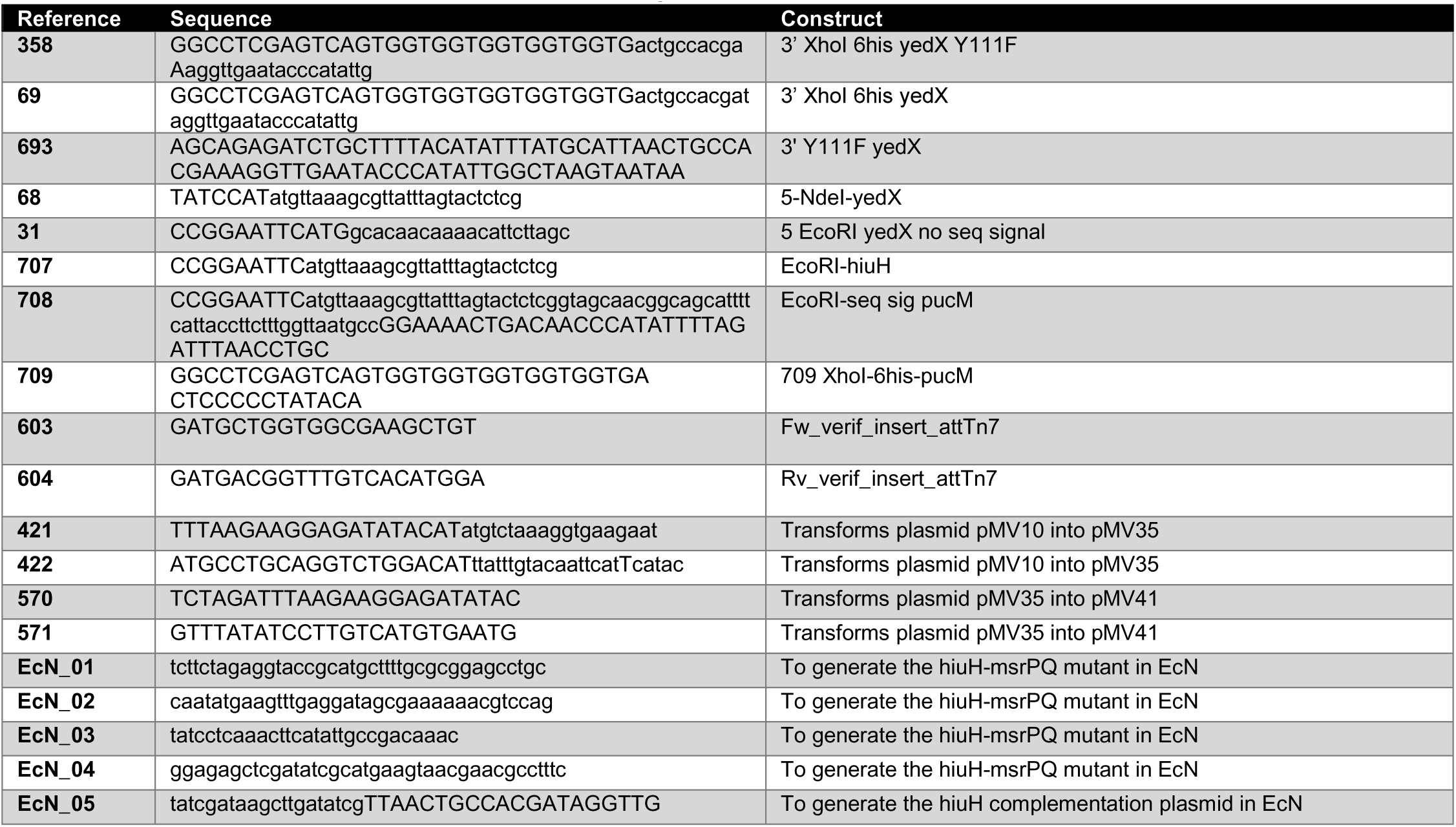
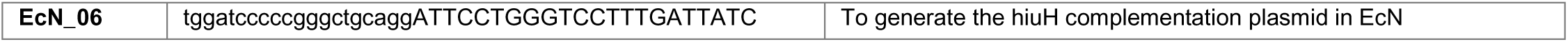
Primers used in this study.

